# Atlas of stress-induced changes in yeast transfer RNA modification levels

**DOI:** 10.64898/2026.08.17.745200

**Authors:** Matea Radešić, Jenni K. Pedor, M. Suleman Qasim, Anna-Emilia Rajaveräjä, Nina Sipari, L. Peter Sarin

**Affiliations:** RNAcious laboratory, Department of Molecular and Integrative Biosciences, Faculty of Biological and Environmental Sciences, University of Helsinki, Helsinki, Finland; School of Pharmacy, Faculty of Health Sciences, University of Eastern Finland, Kuopio, Finland; Doctoral Programme in Molecular and Cellular Systems of Life, University of Helsinki Doctoral School, University of Helsinki, Helsinki, Finland; Doctoral Programme in Multidisciplinary Research in Renewable Natural Resources, University of Helsinki Doctoral School, University of Helsinki, Helsinki, Finland

**Keywords:** tRNA modifications, *Saccharomyces cerevisiae*, epitranscriptomics, long-term stress

## Abstract

Transfer RNA (tRNA) modifications are essential for accurate translation and cellular adaptation to environmental changes. Although short-term modification dynamics are well documented, the impact of prolonged stress exposure on the global tRNA landscape remains largely unexplored. Here, we provide the first systematic profiling of tRNA modifications in *Saccharomyces cerevisiae* following long-term exposure to distinct stress types, including heat, suboptimal pH, oxidative stress (paraquat and diamide), osmotic stress (NaCl and KCl), and genotoxic stress (MMS). Using our broad-range UPLC-MS protocol, we characterized relative nucleoside modification changes across the global tRNA landscape, revealing that long-term stress triggers a global reprogramming of the tRNA epitranscriptome in a stress-specific and time-dependent manner. Remarkably, we identified that pH stress and paraquat induce a near-complete loss of 5-methoxycarbonylmethyl-2-thiouridine (mcm^5^s^2^U_34_) modification, and we observe an increase in the non-thiolated 5-methoxycarbonylmethyl (mcm^5^U_34_) precursor at pH 7. This coupled response is akin to that previously reported for temperature-dependent thiolation deficiency. However, the impact on thiolation is transient in the case of pH stress, but not with paraquat, suggesting two distinct stress-dependent impairment mechanisms of the thiolation pathway. To further integrate our results, we sought to normalize changes in nucleoside modification levels against potential alterations in the tRNA pool. Thus, we performed MarathonRT-based tRNA sequencing and devised the modification deviation (*MD_m_*) index. This established that the observed modification changes occurred independently of tRNA isoacceptor abundance, implying that tRNA modification levels are predominantly affected by other factors. Together, this study provides a comprehensive atlas of tRNA modification dynamics under prolonged stress, addressing a critical gap in our understanding of RNA-based translational control. Furthermore, we present the *MD_m_* index as a robust quantitative framework to decouple the influence of tRNA abundance from global modification signals, providing a necessary metric for the field to interpret epitranscriptomic reprogramming.

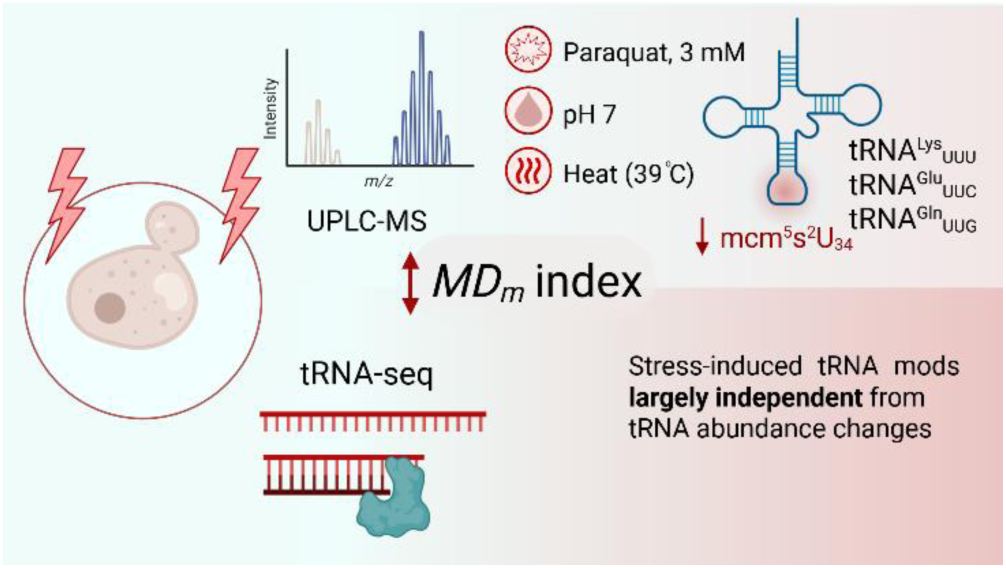

## INTRODUCTION

Transfer RNA (tRNA) is a critical adapter molecule that ensures accurate decoding of the genetic code. This functional flexibility is largely achieved through the plethora of chemical modifications that adorn tRNAs. To date, over 150 different modified ribonucleosides have been identified, and some are found across all domains of life, with tRNAs containing the highest number of modifications and the greatest chemical diversity.^1^ Each tRNA molecule contains 13 biochemical modifications on average, ranging from basic structural modifications such as methylation or acetylation to highly complex chemical alterations.^2^ Modifications occur at specific positions within the tRNA molecule and contribute to its structural stability or decoding capability. For instance, the anticodon stem loop (ASL) is heavily modified, with modifications at position 34 providing decoding flexibility ^3, 4, 5^ whereas modifications at position 37 offering stabilization of codon-anticodon interaction ^6^ and reading frame maintenance ^7^.

The vast chemical diversity of the tRNA epitranscriptome is introduced post-transcriptionally by a repertoire of modifying enzymes.^8, 9^ These processes vary significantly in their biosynthetic complexity. While many “simple” modifications, such as methylations, are typically introduced by the action of a single enzyme, others require the coordinated activity of complex enzymatic pathways.^8, 9^ For example, the biosynthesis of 2-thiouridine (s^2^U) derivatives at wobble position 34 in cytoplasmic tRNA—where the 5-methoxycarbonylmethyl (mcm^5^) precursor is transformed into 5-methoxycarbonylmethyl-2-thiouridine (mcm^5^s^2^U)—relies on a sophisticated multi-enzyme pathway.^10^ This process begins with sulfur activation by the cysteine desulfurase Nfs1^11^, followed by its transfer through the relay protein Tum1 ^12^. The cascade further involves Uba4 and the sulfur carrier Urm1 ^13, 14^, after which the sulfur moiety is finally installed onto the target tRNA by the Ncs2/Ncs6 complex ^10, 12, 13, 15^. Because these modifications are essential for translational fidelity and protein homeostasis, any dysfunction within this cascade can trigger significant modification deficits, that impairs the cell’s ability to adapt to environmental pressures.^3, 13, 15–18^

Yeasts are ubiquitous microorganisms that thrive in different natural habitats, where they face diverse environmental stresses, such as suboptimal temperature, pH fluctuations, high salinity, etc., which require robust adaptive responses.^19–23^ Besides their ecological role, yeasts and particularly baker’s yeast, *Saccharomyces cerevisiae*, is of significant economic relevance in food production, brewing, and biotechnology.^24, 25^ In industrial settings, yeast cultures are frequently subjected to harsh and fluctuating conditions, such as high ethanol concentrations, osmotic shifts, and nutrient limitation. ^26, 27^ Consequently, understanding cellular resilience and achieving optimized resource utilization is vital for enhancing biomanufacturing efficiency and product yields. ^28^

Owing to its well-characterized genome and widely available tools for genetic manipulation, *S. cerevisiae* has been established as an appropriate model organism for studying the tRNA epitranscriptome. ^29, 30^ Previous studies in yeast have established that tRNA modifications are dynamic and rapidly adjust to sudden biotic and abiotic changes. ^16, 31–36^ Most of the seminal work has focused on short-term stress exposure, demonstrating how rapid shifts in anticodon stem loop (ASL) modification levels trigger codon-biased translation of specific stress-response proteins to enhance cell survival. ^4, 31, 32, 34, 35, 37, 38^

While short-term tRNA modification dynamics are relatively well understood, the effects of prolonged stress exposure on the global tRNA modification landscape remain largely unexplored. This gap is critical because long-term stress conditions in yeast are physiologically relevant in industrial fermentation, pathogenic persistence, and environmental survival.^26, 39–41^ Revealing how tRNA modification profiles evolve under extended stress provides further understanding of translational control and opens opportunities for applied biotechnology. If specific modification signatures can be reproducibly induced, they could be leveraged to engineer stress-resilient strains or optimize protein synthesis under defined conditions. Such control over tRNA modification profiles could enable tailored translation programs for biomanufacturing, improving yields of stress-sensitive products or enhancing robustness in large-scale processes.

Here, we present the first systematic profiling of tRNA modifications in yeast after long-term exposure to seven distinct stressors. Using our broad-range UPLC-MS protocol ^42^, we characterized changes in tRNA modifications and identified stress-specific signatures. Notably, a complete loss of thiolation was observed at elevated temperatures, and a partial loss occurred after exposure to paraquat and pH stress in isoacceptors tRNA^Lys^(UUU), tRNA^Glu^(UUC), tRNA^Gln^(UUG). While the temperature-sensitivity of mcm^5^s^2^U is a well-documented phenomenon ^16, 35, 41, 43, 44^, this is the first time that a similar decrease in mcm^5^s^2^U levels is observed for other stressors. We compared tRNA expression levels under selected stress conditions, showing that the levels of some isoacceptors fluctuate when stress is applied. Since fluctuations in tRNA isoacceptor abundance directly undermine the quantitative assessment of tRNA modification levels, we sought to establish a metric that accounts for the tRNA modification signal against its prevalence in the tRNA pool. To this end, we developed a modification deviation (*MD_m_*) index that weighs the contribution of individual isodecoders according to their baseline abundance and modification frequency. By providing a comprehensive overview of long-term, stress-induced reprogramming of tRNA modifications, this study addresses a critical gap in our understanding of translational control under sustained stress.

## MATERIALS AND METHODS

### Strains and media

*Saccharomyces cerevisiae* strain BY4741 was used for all experiments in this study. *S. cerevisiae* cultures were plated from glycerol stock to Yeast Extract Peptone Dextrose (YPD) agar plates containing 1% yeast extract (Neogen), 2% peptone (Gibco), 2% glucose (VWR Chemicals), and 2% agar (Neogen), and grown at 30 °C for 48 h. Liquid YPD medium contained 1% yeast extract (Gibco), 2% peptone (Gibco), and 2% glucose (Acros Organics). Liquid cultures were grown in YPD broth at 30 °C, unless otherwise stated, for 12 h or 24 h. Viable counts (CFU/ml) were determined by plating serial dilutions on YPD agar.

### Cytotoxicity screening

Stress conditions were induced either by incubating the cultures at elevated temperatures (heat stress) or by supplementing the YPD medium with: a) H_3_PO_4_ (Fisher Scientific) for acidic pH stress, b) KOH (Fisher Chemical) in combination with 100 mM KH_2_PO_4_ (Arcos Organics) for neutral and alkaline pH stress c) methyl methanesulfonate, MMS (Acros Organics) for genotoxic stress, d) KCl (VWR Chemicals) and NaCl (Fisher Scientific) for osmotic stress, or e) diamide (MP Biomedicals) and paraquat (Aldrich) for oxidative stress (Table S1). Preliminary cytotoxicity screening was conducted by cultivating 200 µL of *S. cerevisiae* cultures in 100-well Honeycomb microplates (Oy Growth Curves Ab Ltd). Cultures were grown from a starting OD_600_ of ∼0.2. Stress was applied to starter cultures and continued throughout the whole assay. Growth was continuously monitored by measuring the optical density (OD_600_) every 10 min over a 24 h period using a Bioscreen C (Oy Growth Curves Ab Ltd) microplate photometer. Cell survival was calculated as the percentage of viable cells in stressed cultures relative to the untreated control at each corresponding timepoint, determined by counting colony-forming units (CFU/mL) after plating harvested cells on YPD agar. Selected stressor doses were further verified in 50 mL YPD liquid cultures, with starting OD_600_ ∼0.2. Growth in flasks was monitored every hour for 12 h, and after 24 h by measuring OD_600_ using a spectrophotometer (Eppendorf). Cells were harvested at 6 h, 12 h, and 24 h for determining their viable counts. The specific growth rate (µ, h^-1^) during the exponential growth phase was calculated as:

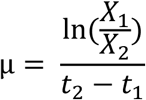

where *X*_1_ and *X*_2_ represent average OD_600_ measurements at timepoints *t*_1_ and *t*_2_ taken during the exponential phase.

### Total RNA & tRNA isolation

*S. cerevisiae* was cultured in 100 mL YPD at 30 °C (except for heat stress) with constant aeration (180 rpm) in the presence of chemical stressors or at elevated temperature (as indicated in Fig. 1C). Total RNA was isolated as previously described.^16, 42^ Briefly, cells were harvested after 12 h and 24 h by pelleting at 4,000× *g* for 10 min at 4 °C. Pellets were washed with 1× phosphate-buffered saline, then resuspended in 1 vol of 0.9 % sodium chloride (NaCl) solution, to which 1 vol of acidic phenol, pH 4.3, (Carl Roth), and 0.2 vol of bromochloropropane (Acros Organics) were added. Glass beads (0.1 and 0.5 mm in a 1:2 ratio) were added to the mixture, and samples were vortexed at maximum speed for 5 min. The mixture was centrifuged at 10,000× *g* for 15 min at room temperature. The aqueous phase was transferred to a new tube containing 0.5 vol of acidic phenol and 0.1 vol of BCP, vortexed vigorously, and recentrifuged at 10,000× *g* for 10 min at room temperature. RNA was precipitated from the resulting aqueous phase using 2.5 vol of 99.6 % ethanol at −20 °C overnight. Total RNA was pelleted at 10,000× *g* for 30 min at 4 °C, washed three times with 80 % ethanol and centrifuged at 10,000× *g* for 20 min at 4 °C. The RNA pellet was air-dried, resuspended in RNase-free double-distilled water (ddH_2_O) and stored at −70 °C.

**Figure 1.**
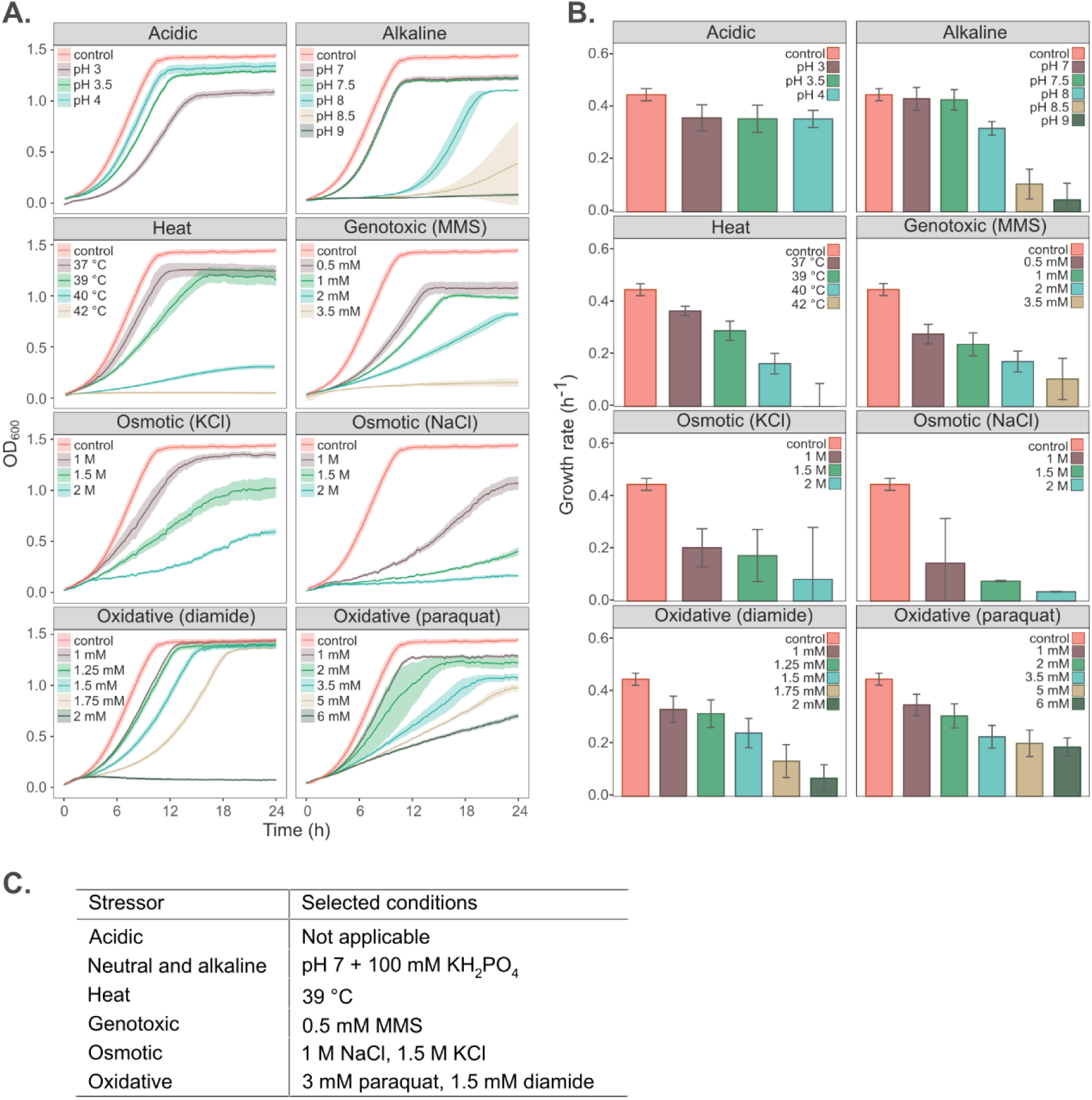
Initial screening of *S. cerevisiae* growth dynamics at different stress conditions. (A) Growth curves following exposure to various stressors and doses. Each panel represents a single stressor and each line color represents a different dose intensity (from weakest to strongest; brown, green, dark cyan, beige, and dark green): acidic stress (pH 3–4), alkaline stress (pH 7–9), genotoxic stress (MMS, 0.5–3.5 mM), heat stress (37– 42 °C), osmotic stress (KCl and NaCl, 1–2 M), and oxidative stress (diamide, 1–2 mM; paraquat, 1–6 mM). The control culture (YPD, 30 °C) is shown as a red line. Ribbon shading indicates standard deviation (SD). (B) Specific growth rates (h^-1^) are calculated based on the early exponential phase (derived from growth curves in A). Data represents the mean ±SD from independent biological replicates (n = 3). C) Selected conditions yielding a moderate impact (60–80% reduction in survival) on the *S. cerevisiae* growth rate.

tRNA was isolated as previously described with minor adjustments ^45^. Briefly, 100 µg of total RNA was mixed with 250 µL of Long RNA Binding Buffer (5 M guanidinium thiocyanate, 50 mM Tris-HCl, pH 7.0) and 190 µL of 100 % ethylene glycol diacetate (EGDA). The mixture was loaded onto a NucleoSpin RNA Column (Macherey-Nagel) placed into a 2 mL tube and centrifuged at 11,000× *g* for 1 min at room temperature. The flowthrough was collected, mixed with 675 µL of Short RNA Binding Buffer (80 % EGDA, 1 M guanidinium thiocyanate, 10 mM Tris pH 7.0), vortexed thoroughly, and incubated for 1 min at room temperature. The mixture was then loaded onto a new NucleoSpin RNA Column, spun at 11,000× *g* for 1 min, and washed with 500 µL of Chaotropic Wash Buffer (1.6 M guanidinium thiocyanate, 66 % ethanol, 20 mM Tris-HCl, pH 7.0) followed by 500 µL of Ethanol Wash Buffer (10 mM Tris-HCl pH 7.5, 20 mM NaCl, 80 % ethanol). The empty column was spun for 2 min at 11,000× *g* to remove residual wash buffer. Bulk tRNA was eluted with 50 µL of RNase-free ddH_2_O and stored at −70 °C. The quality and purity of the isolated tRNA was analyzed by 10 % urea-polyacrylamide (urea-PAA) gel electrophoresis. The concentration was measured on Nanodrop (Thermo Fisher Scientific).

### Monoribonucleoside analysis by LC-MS

The digestion of ribonucleosides proceeded as previously described.^16, 42^ Briefly, 5-10 µg of tRNA was digested with Nuclease P1 from *Penicillium citrinum* (Sigma-Aldrich) and Fast AP Thermosensitive Alkaline Phosphatase (Thermo Scientific) at 37 °C in the presence of ZnCl_2_ (Alfa Aesar) and sodium acetate, pH 5.3. The reaction was terminated by adding trifluoroacetic acid (Acros Organics) to a final concentration of 1.0 %. Ribonucleosides were purified with Thermo Hypersep™ SpinTip Hypercarb™ (Thermo Fisher Scientific), dried in a SpeedVac concentrator (Thermo Fisher Scientific), and resuspended in 5 mM ammonium formate, pH 5.3. For data normalization, 1,3-dimethylpseudouridine (m^1,3^Ψ) was added in each sample as an internal standard (0.1 µg of m^1,3^Ψ per 10 µg of bulk tRNA). Next, ribonucleoside standards and biological samples were analyzed with Waters Acquity UPLC system coupled to a Waters Synapt G2-Si HDMS mass spectrometer equipped with an electrospray ionization (ESI) source. Analyses were performed in positive ion mode over a mass range of 90–800 m/z. Analytical parameters were set as previously described.^42^

### tRNA separation on APM-urea-PAA gel and Northern blot

The analysis was essentially performed as previously described.^45^ Briefly, 500 ng of tRNA was separated on urea-polyacrylamide gel containing 5 µg/mL [(N-Acryloylamino)phenyl]mercuric chloride (APM), (Fisher Scientific). Afterwards, the gel was soaked for 1 h in 10 mM dithiothreitol (DTT), (Fisher BioReagents) dissolved in 0.5x TBE, following the transfer on an Amersham™ Hybond-N+ nylon membrane at 300 mA, 24 V for 1 h. Next, the transferred tRNA was UV-crosslinked onto the membrane, which was then prehybridized for 1 h and hybridized overnight with biotinylated DNA probes (Table S4) at 55 °C. The next day, washing and blocking of the membrane were performed as described, following the chemiluminescent detection.

### tRNA sequencing

tRNA sequencing was performed as described previously.^46, 47^ Briefly, total RNA was deacylated and dephosphorylated, following the enrichment of mature tRNAs on the silica-based spin column.^45^ Sequencing libraries are prepared according to the mim-tRNAseq workflow ^48, 49^, involving 3’ adapter ligation and reverse transcription at 42 °C for 16 hours using 0.5 U of MRT-CBD per 100 ng of tRNA. After cDNA circularization and library amplification, samples are sequenced on an Illumina platform and analyzed using the mim-tRNAseq toolkit to compare tRNA abundance in different conditions.^48, 49^

### Quantitative assessment of tRNA modification regulation by modification deviation index

The modification deviation index was created to quantify the dependency of tRNA modification levels changes linked to the abundance of modification-carrying tRNAs, and it is calculated as:

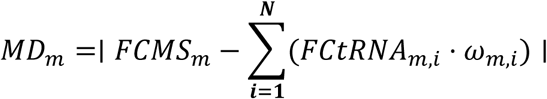

where *MD_m_* is the modification deviation index, *FCMS_m_* is the centered observed fold change of modification *m* measured by UPLC-MS between the stressed condition and control, and *FCtRNA_m_*_,*i*_ is the centered fold change of tRNA isodecoder *i* obtained from tRNA-seq analysis. The term *∑^N^_i=1_* (*FCtRNA_m_*_,*i*_ ⋅ *ω_m_*_,*i*_) represents the expected abundance-driven change in the modification signal, calculated as the weighted sum of the fold changes of all *N* tRNA isodecoders known to carry modification *m*. The weighting factor *ω_m_*_,*i*_ describes the fractional contribution of isodecoder *i* to the total cellular abundance of modification *m* and was calculated as:

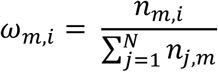

where *n_m_*_,*i*_ represents the number of annotated modification sites *m* per isodecoder *i*, and *∑^N^_j=1_ n_j_*_,*m*_ represents the total number of sites for that modification across the total number of isodecoders *N* annotated in tModBase. Weighting factors were determined for 47 cytoplasmic and mitochondrial isodecoders (Supplementary File S1, S2). An *MD_m_* index threshold of > 0.2 was empirically set to denote tRNA modification level changes that are independent from fluctuations in tRNA abundance.

### Data analysis

Peak identification from mass spectrometry data was performed using Mzmine 4.5.37 ^50^ based on a custom lookup list of ribonucleosides consisting of previously reported [M+H]^+^ and product ion masses available in MODOMICS ^1^, as well as retention time obtained from available ribonucleoside standards.^42^ Normalization was performed using the internal spike-in standard m^1,3^Ψ. Ribonucleoside abundances were semi-quantitatively characterized by their relative change, where normalized peak intensities from stressed samples were compared to their respective controls and expressed as stress/control fold changes. Statistical analysis was performed using unpaired two-sample *t*-tests, with error bars representing the standard deviation from 5 biological replicates, unless otherwise stated. Pearson’s correlations were computed in R (version 2025.05.1). Microsoft Excel was used for supplementary calculations and *MD_m_* index calculations. Sequencing data analysis was performed as previously described^46, 47^. Sequencing was performed on 5 biological replicates, unless otherwise stated. Figures and tables were created and visualized in R (version 2025.05.1), Microsoft Excel, and Inkscape (version 1.4.2).

## RESULTS

### Environmental and chemical stressors modulate growth dynamics of S. cerevisiae

To investigate how tRNA modification profiles respond to environmental stress, *S. cerevisiae* BY4741 was initially exposed to eight distinct stress conditions representing different mechanisms of toxicity (Table S1). A preliminary small-scale cytotoxicity screening was performed to identify stress intensities that produced comparable phenotypic endpoints across conditions. We observed that all stressors induced a dose- and stress-dependent reduction in *S. cerevisiae* growth (Fig. 1). Most stresses inhibited proliferation by extending the lag phase (Fig. 1A) and progressively reducing the growth rate (Fig. 1B) as the concentration or intensity of the stressor increased. Acidic stress slightly impaired growth, with the strongest inhibition observed at pH 3, the lowest pH tested. *S. cerevisiae* is known to grow optimally under mildly acidic conditions (pH 4–6) and actively acidifies its surroundings ^51^, which accounts for why the pH range (pH 3–4) used here does not severely inhibit its growth. In contrast, a clear growth inhibition was observed in alkaline conditions at pH 8 or higher, resulting in a growth delay of approximately 9 h before the culture enters exponential growth phase (Fig. 1A). Osmotic stress caused by high salt concentrations also inhibited growth, with NaCl exerting a stronger effect than KCl at equivalent concentrations. Oxidative stress induced by paraquat and diamide produced a pronounced growth inhibition, with diamide suddenly completely abolishing growth at concentrations ≥2 mM. Genotoxic stress by methyl methanesulfonate (MMS) led to a dose-dependent decrease in growth, with complete inhibition at 3.5 mM, whereas elevated temperatures (37–42 °C) were similarly detrimental, and growth was completely inhibited at >40 °C. Therefore, *S. cerevisiae* exhibits sensitivity to a broad range of environmental and chemical stressors, and the specific type of stress strongly influenced the growth dynamics, and in all cases, growth was suppressed in a dose-dependent manner.

### Stress survival patterns highlight different adaptation responses

To validate the growth trends from the initial Bioscreen screening (Fig. 1B), we selected stress conditions that moderately reduced the growth rate (∼60–80 % relative to the control; Fig. 1C). We proceeded with scaling-up these exposures to larger liquid cultures in flasks to independently verify growth inhibition and assess survival over time (Fig. 2). Overall, exposure to the stressors caused a substantial decline in viable counts, with only a few conditions allowing partial recovery. Growth under pH stress, elevated temperatures, and exposure to MMS, KCl, or paraquat, triggered a continuous decline in survival with negligible signs of recovery (i.e., an increase in the percentage of viable cells at a later timepoint compared to an earlier one). Among these, pH stress resulted in the lowest viable counts at both timepoints (Fig. 2B). In contrast, diamide exposure caused an initial drop in viable counts at 6 h, followed by a sharp recovery at 12 h (72.8 % survival), before declining again at 24 h (Fig. 2A, B). Osmotic stress induced by NaCl led to decreased survival, with partial recovery observed only at 24 h, with 57.3 % of the cells surviving the stress compared to 24.0 % at 12 h (Fig. 2A, B).

**Figure 2.**
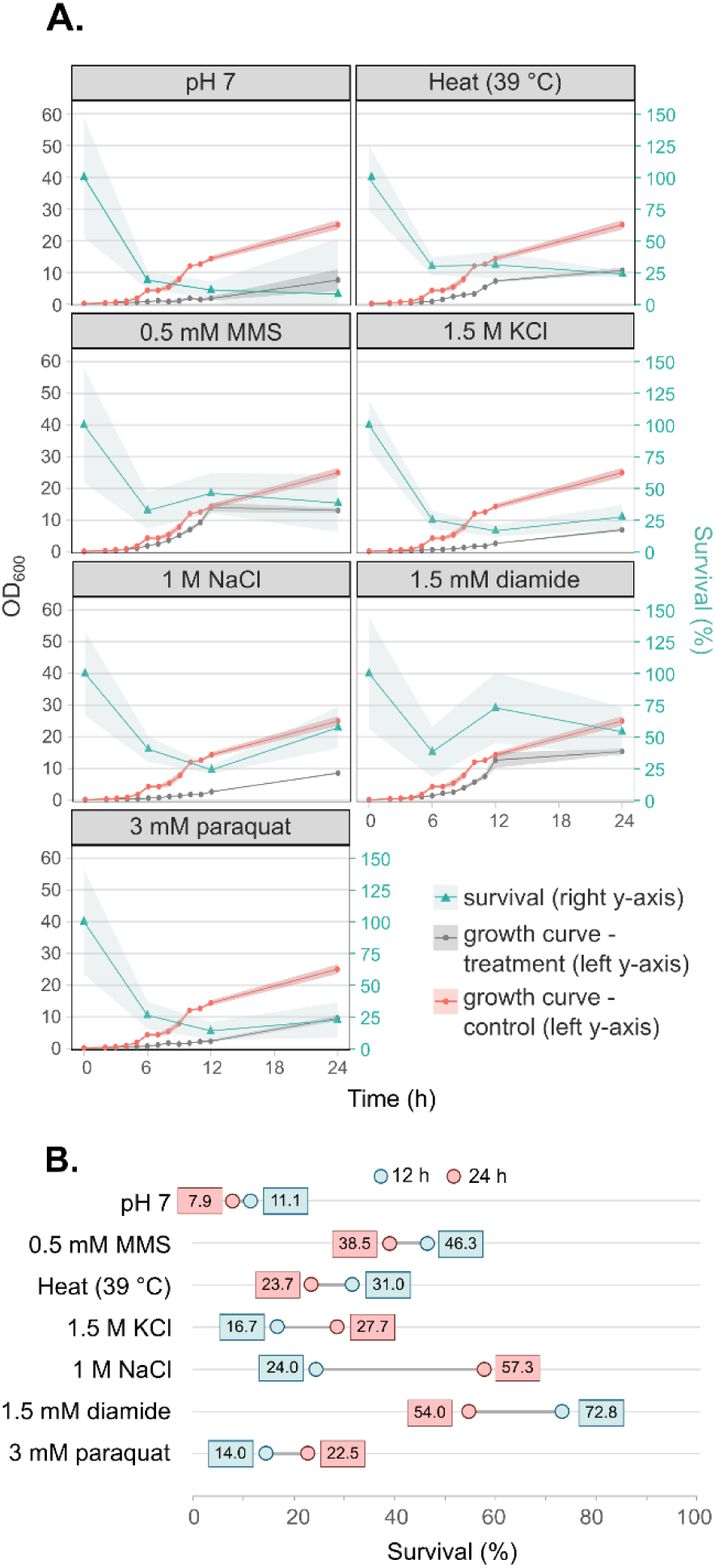
Growth and survival of *S. cerevisiae* under different stress conditions. (A) Growth curves (OD_600_, left y-axis, black) and survival percentages (right y-axis, cyan) are shown for *S. cerevisiae* continuously exposed to various stressors: pH 7, 0.5 mM MMS, heat (39 °C), 1.5 M KCl, 1 M NaCl, 1.5 mM diamide, and 3 mM paraquat. OD_600_ was measured hourly for 12 h and again at 24 h with a spectrophotometer. Circles indicate individual measurement points. For each condition, the control culture growth curve is represented in red, and the stressed culture growth curve in grey, with shaded areas indicating standard deviation. Survival rates are represented by cyan lines with triangle markers representing sampling points at 0, 6, 12 and 24 h. Viable counts were assessed by plating and survival was calculated by comparing CFU/mL of stressed cells to the control. Data represent the mean ± SD from independent biological replicates (n = 3). (B) Comparative dumbbell plot of survival percentages at 12 h (blue) and 24 h (red) derived from A. Numeric labels indicate mean survival values. Data represents the mean from independent biological replicates (n = 3).

### Continued stress leads to global reprogramming of tRNA modifications

To profile global changes in tRNA modifications in long-term, steady-state stressed cultures of *S. cerevisiae*, we performed semiquantitative tRNA modification analysis using our broad-range C18-UPLC-MS approach capable of detecting a wide spectrum of modified ribonucleosides from complex biological samples.^42^ In total, 22 modifications were detected (Fig. 3A, Table S2); N6,N6-dimethyladenosine (m^6,6^A), 5-carbamoylmethyluridine (ncm^5^U) and N4-acetylcytidine (ac^4^C) were identified, but their signals were insufficient for reliable relative quantification, as they fell below the 5:1 signal-to-noise threshold required for peak analysis (Fig. S2). Moreover, 2’-O-methylcytidine (Cm), mcm^5^s^2^U, and occasionally mcm^5^U and 1-methylinosine (m^1^I), exhibited lower signal intensity, but above the threshold of signal-to-noise level (Fig. S2).

**Figure 3.**
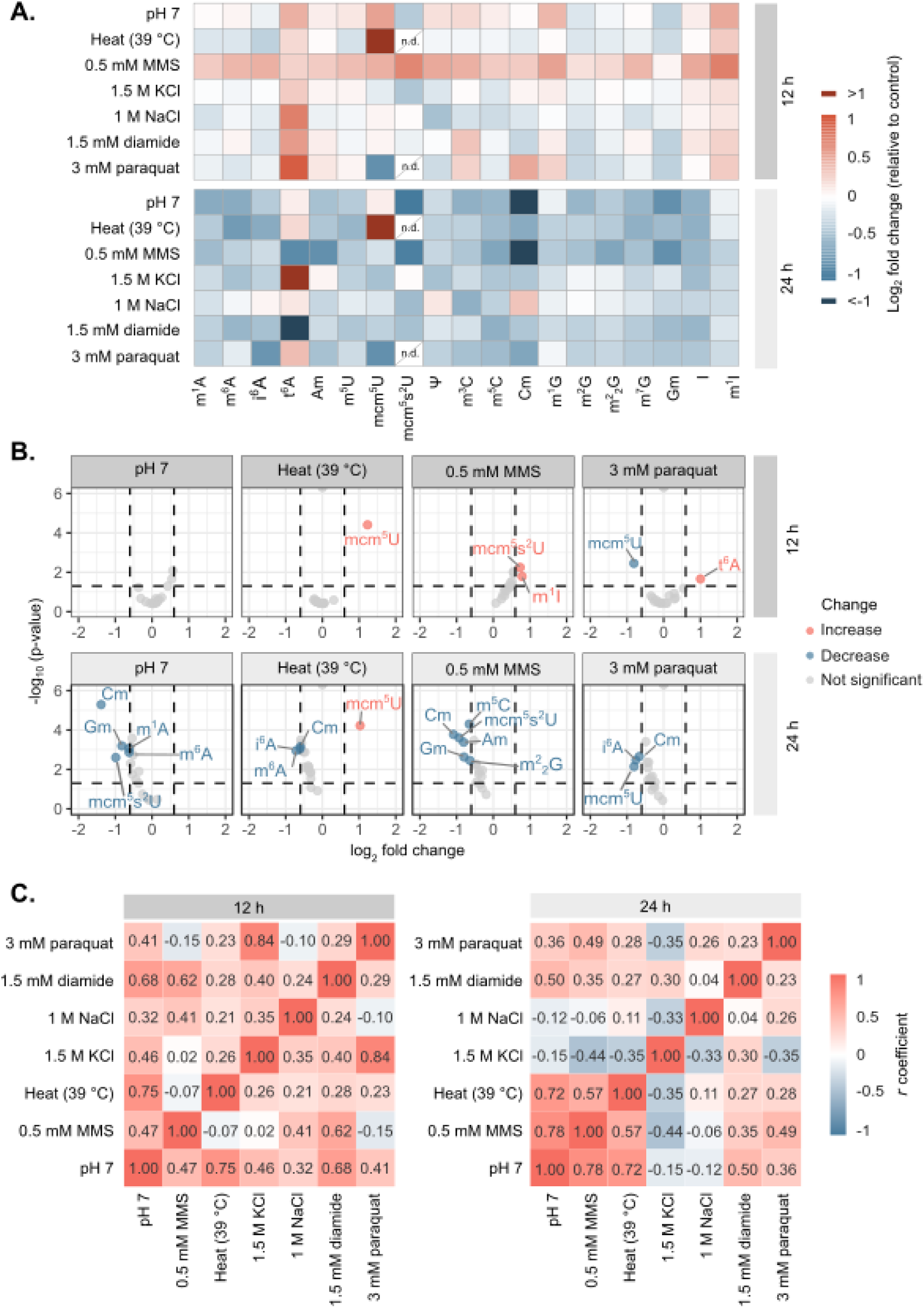
Changes in tRNA modification levels under stress conditions. (A) Heatmap visualization of tRNA modification levels in *S. cerevisiae.* Each column represents a distinct tRNA modification, and each row corresponds to a stress condition. Color intensity reflects the magnitude and direction of change (red – increased levels compared to the control, blue – decreased levels compared to the control, n.d. – not detected. Data are expressed as log_2_ fold change relative to the control, representing the average normalized MS peak height signals from biological replicates (n = 5) analyzed by UPLC-MS, except where n = 4 (paraquat, KCl exposure at 12 h; paraquat, MMS exposure at 24 h) and n = 3 (NaCl exposure at 12 h). Normalization was performed against an internal standard (non-natural modification m^1,3^Ψ). Abbreviations for tRNA modifications follow the short names used in MODOMICS.^1^ (B) Volcano plots show the distribution of tRNA modifications in response to selected stressors. Each point represents a tRNA modification, plotted by log_2_ fold change (x-axis) and statistical significance as -log_10_ of p-value (y-axis). P-values were calculated using an unpaired two-samples *t*-test. Points meeting significance criteria (|log_2_ fold| > 0.6 and p < 0.05) are colored in coral red (increased levels compared to the control) and blue (decreased levels compared to the control). Non-significant points are shown in grey. (C) Heatmap showing pairwise correlation of changes in tRNA modifications between different stress conditions applied to samples at two timepoints, 12 h and 24 h. Each cell shows the Pearson’s *r* correlation coefficient, with positive and negative values colored in shades of red and blue, respectively. Correlation is calculated from data in A.

As expected, we observed a global stress-dependent decrease for most modification levels (Fig. 3).^16, 33^ Generally, a greater proportion of modifications exhibited significant changes after 24 h compared to 12 h (Fig. 3B). To ensure that these time-dependent changes were accurately represented, all ribonucleoside abundances were compared relatively to their equivalent control (i.e., stressed cultures at 12 h samples were normalized to the control at 12 h). However, it is important to note that the control samples themselves differ significantly between the two timepoints (Fig. S3), reflecting the cells’ continuous growth and transition into a different metabolic state at later timepoints, characterized by high cell density and early stationary phase physiology. Notably, Cm levels were consistently lower in all stress samples at 24 h (Fig. 3A, B); yet, when comparing controls, Cm was approximately two-fold higher at 24 h than at 12 h (Fig. S3). Given the low standard deviation observed, this signal is likely a biological reflection of the transition to high-cell-density growth rather than a technical artifact. However, because the signal intensity for Cm is relatively low and close to the 5:1 signal-to-noise threshold required for peak analysis, it should be interpreted with caution.

### 2-thiolation is sensitive to prolonged exposure to specific stressors

Upon closer inspection of the stress-specific signatures at 12 h, we found that after exposure to MMS, there was a slight enrichment of most modifications, with mcm^5^s^2^U and m^1^I being significantly increased. Under heat stress, mcm^5^U showed a strong increase, whereas its thiolated form, mcm^5^s^2^U, could not be detected. Paraquat exposure led to a small but significant decrease in mcm^5^U and, similar to heat stress, mcm^5^s^2^U was below the detection limit. Notably, growth at pH 7 displayed a comparable pattern, characterized by a modest increase in mcm^5^U and a decrease in mcm^5^s^2^U, although these alterations were less pronounced than in heat stress (Fig. 3A, B).

Conversely, at the 24 h time point, we noticed a strong increase in mcm^5^U levels for the heat stressed samples, while Cm and m^6^A were decreased, and mcm^5^s^2^U was not detected (Fig. 3A, B). MMS exposure caused a significant decrease in Cm, Gm, Am, m^2^_2_G, mcm^5^s^2^U, and m^5^C levels. Paraquat exposure affected Cm and mcm^5^U, while mcm^5^s^2^U was—as in the previous timepoint—below the detection limit, whereas neutral pH caused Cm, Gm, m^1^A, m^6^A and mcm^5^s^2^U levels to decrease. In contrast, tRNA modification profiles remained largely unaltered under osmotic stress (KCl, NaCl) and diamide exposure, with only a single modification showing a significant shift after 24 h (Fig. S4). Interestingly, many of the significantly altered modifications are found in the anticodon loop, including U_34_ wobble modifications mcm^5^s^2^U and mcm^5^U. Such changes may alter translational flexibility, enabling the synthesis of diverse proteins under suboptimal environmental conditions ^3, 4, 18,^ _32, 34._

A common stress-induced signature observed across three stress exposures (heat stress, oxidative stress caused by paraquat, and suboptimal pH) was the complete or partial loss of detectable thiolation signal for the mcm^5^s^2^U modification in the UPLC-MS analysis (Fig. 3A). While heat-induced thiolation loss has been previously reported ^16, 35, 41, 43, 44^, the reductions observed under paraquat and suboptimal pH exposure represent novel findings. However, the overall mcm^5^s^2^U signal produced by the UPLC-MS analysis was of relatively low intensity, which may limit the precision of its characterization. To independently validate these findings, we performed APM-affinity gel electrophoresis and Northern blotting ^45^, and we evaluated the thiolation status of all three cytoplasmic isoacceptors known to carry mcm^5^s^2^U: tRNA^Glu^(UUC), tRNA^Lys^(UUU), and tRNA^Gln^(UUG) (Fig. S5).

The Northern blot analysis confirmed a significant reduction in the thiolated tRNA species across the tested conditions (Fig. S5A, B), consistent with the UPLC-MS analysis (Fig. 3A). Unsurprisingly, control samples maintained stable levels of highly thiolated tRNAs, with only ∼10 % of non-thiolated species detected for tRNA^Glu^(UUC). Exposure to pH 7 resulted in a significant reduction of detectable thiolated tRNA species, however, we observed a partial recovery of the signal at 24 h relative to the 12 h timepoint, indicating a time-dependent adaptive response. Interestingly, this major change occurred despite minimal recovery in OD_600_ or viable counts between the tested interval (Fig. 2), potentially showing that cells are still actively modulating translation at high-stress conditions. As expected, heat stress induced a loss of thiolation signal across all timepoints and tRNA probes tested. Although some traces migrating at the expected position for thiolated tRNA^Gln^(UUG) were visible (Fig. S5A), the signals remained below the threshold for reliable densitometric quantification. Paraquat exposure led to a noticeable decrease in the thiolated tRNA species (Fig. S5A, B), confirming a partial decrease in thiolation. This finding clarifies the UPLC-MS results, which suggested a near-complete loss, as the modification signal fell below the instrument’s detection limit (Fig. 3A). While the magnitude of paraquat’s effect was highly isoacceptor-specific, it remained stable across both timepoints, distinguishing it from the time-dependent thiolation response observed under pH 7 stress. Together, these results highlight the effect of time, stressor, and isoacceptor on mcm^5^s^2^U dynamics and revealed that paraquat and pH-induced stress also cause decreased thiolation, which to the best of our knowledge represents a novel finding.

### Heat, paraquat, and pH stress show stable tRNA modification profiles over time

To assess the relationship between tRNA modification patterns under different stress conditions and across time, we performed correlation analysis using normalized fold-change values (i.e. exposure /control) (Fig. 3C, Fig. S6A), consistent with the previously used values (Fig. 3A, B). When comparing the same condition over time, paraquat, elevated temperature, and pH 7, the profiles exhibited strong positive correlations, indicating that these stresses maintain a similar modification profile over time (Fig. S6A). This observation aligns with the UPLC-MS results (Fig. 3A) and Northern blots (Fig. S5), where heat stress consistently resulted in loss of thiolation and a pronounced increase in mcm^5^U at both timepoints. Similarly, paraquat exposure initially led to a decrease in mcm^5^s^2^U at both timepoints, keeping the thiolation levels low over the duration of the experiment. Additionally, growth at pH 7 followed a comparable trend across both timepoints. In contrast, other stress conditions displayed a more random distribution of modifications, suggesting no clear correlation.

When comparing different stress conditions at the same timepoint, distinct correlation patterns emerge (Fig. 3C). At 12 h, we observed the following conditions to highly correlate: paraquat with KCl, heat stress with pH 7, diamide with pH 7, and MMS with diamide. Surprisingly, heat stress and paraquat exposure did not exhibit strong correlation, regardless of the similar effect those stressors exhibited to mcm^5^s^2^U (Fig. 3A, Fig. S5A, B). At 24 h, heat stress and pH 7 remained highly correlated, with an additional strong correlation between pH 7 and MMS. These results suggest that most stress conditions induce unique modification signatures rather than converging on a common pattern. Moreover, it suggests that pH 7 and elevated temperature may activate a similar stress response pathway.

### Global remodeling of tRNA isoacceptor abundance under long-term stress

The composition of cellular tRNA pools is critical for efficient mRNA decoding and maintaining proteome integrity. The abundance, charging, and modification status of individual tRNA species can differ in distinct cellular environments.^49^ To quantitatively profile the cytosolic tRNA pool in *S. cerevisiae* under long-term stress conditions, we performed tRNA sequencing using Marathon RT (MRT), a highly processive group-II–intron reverse transcriptase.^46, 47^ This enzyme exhibits properties that reduce RT drop-offs at modified sites and favor single-nucleotide substitution signatures over truncations, enabling quantitative, position-resolved detection of multiple tRNA modifications in the same run.^52^ This approach allowed us to assess stress-dependent changes in nuclear tRNA isoacceptors’ and isodecoders’ abundance across a variety of stress types and duration.

tRNA sequencing was performed on samples derived from cultures exposed to four selected conditions (pH 7, heat, MMS and paraquat), which displayed multiple significant changes in tRNA modifications (Fig. 3B). The RNA isolated from the same samples used for UPLC-MS analysis was processed for tRNA sequencing. The results revealed significant changes of the nuclear tRNA pool under tested long-term stress conditions (Fig. 4A). Suboptimal pH was the only condition in which the number of significantly altered tRNA isoacceptors increased between 12 h and 24 h, indicating progressive adaptation. In contrast, heat stress and paraquat exposure showed the same number of significantly up- or downregulated tRNA isoacceptors at both timepoints, although their identity differed over time. The most unexpected pattern was observed under MMS exposure: while nine tRNA isoacceptors were differentially expressed at 12 h, none remained significantly altered at 24 h. To ensure that this temporal variation was not due to the use of separate control samples (i.e., stressed samples were normalized to corresponding timepoint control), we compared tRNA abundances in controls across both timepoints and found no significant differences (Fig. S8), confirming that the MMS-induced change was not affected by control variability.

**Figure 4.**
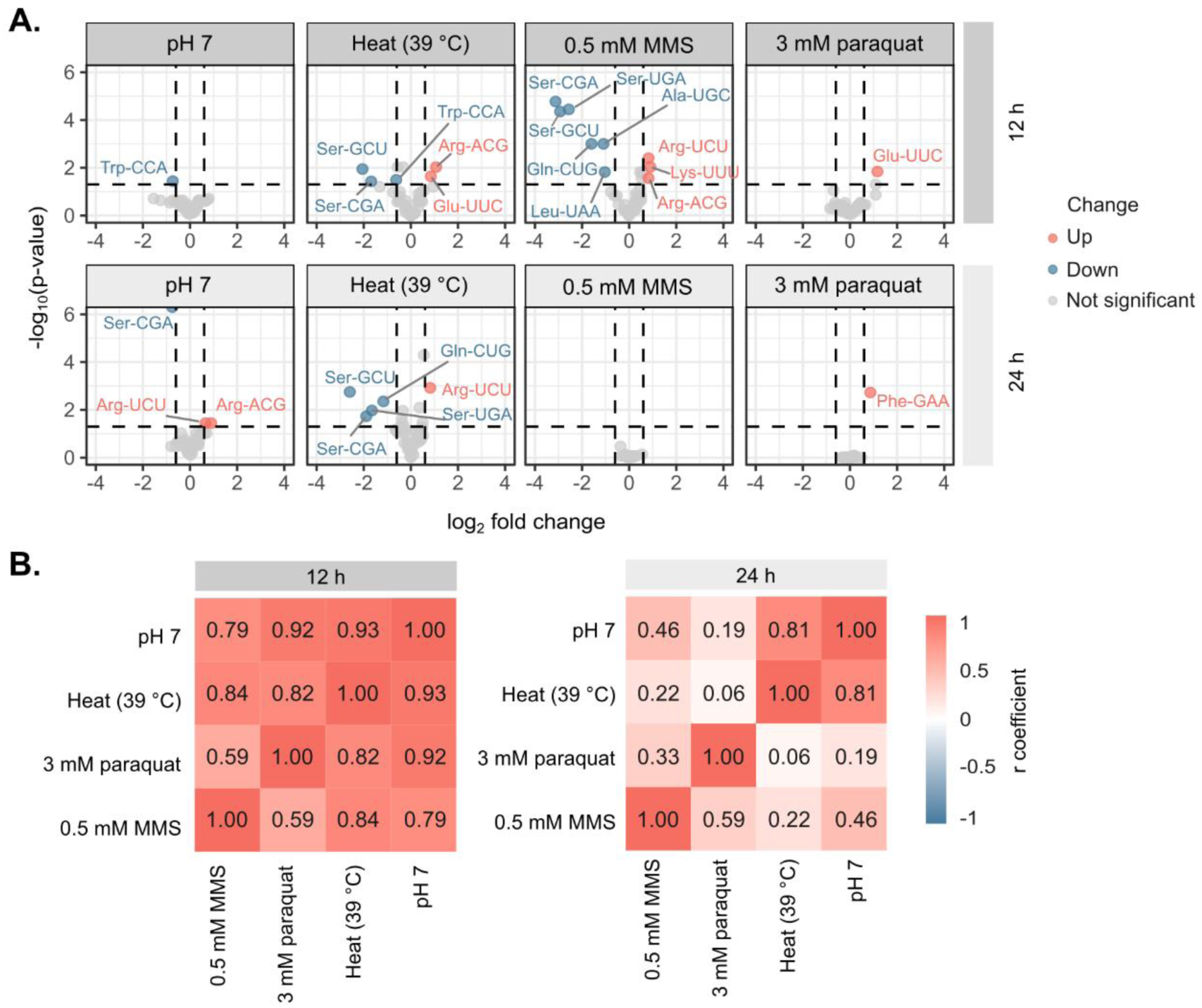
Changes in tRNA isoacceptor expression under selected stress conditions. (A) Volcano plots show the distribution of tRNA isoacceptors in response to each stress exposure for 12 h and 24 h. Each point represents a tRNA isoacceptor, plotted by log_2_ fold change (x-axis) and statistical significance as -log_10_ of p-value (y-axis). Points meeting significance criteria (|log_2_ fold| > 0.6 and p < 0.05) are colored in coral red (upregulation) or blue (downregulation). Dashed vertical lines mark log_2_ fold change thresholds (± 0.6), and the dashed horizontal line marks the significance threshold (p = 0.05). Data are derived from tRNA-seq analysis across independent biological replicates (n = 5), except for samples where n = 4 (control at 12 h) and n = 3 (paraquat at 12 h). (B) Heatmap showing pairwise correlation of changes in tRNA abundance between different stress conditions applied to samples at two timepoints, 12 h and 24 h. Each cell shows the Pearson’s *r* correlation coefficient, with positive and negative values colored in shades of red and blue, respectively. Correlation is calculated from data in A.

### Stress-induced tRNA modification changes are independent from isoacceptor expression

Since the tRNA pool undergoes stress-induced fluctuations, this implies that the observed tRNA modification profiles represent either changes to the modification level itself, to the abundance of the tRNA molecule that carries the modification, or both factors combined.^53, 54^ To account for this, we established a modification deviation (*MD_m_*) index, defined as the absolute difference between the observed fold change in the modification levels and their expected change based on the weighted abundance of their tRNA carriers. Analysis across 47 isodecoders revealed most of the modification that exhibited significant change (Fig. 3B) showed an *MD_m_* score > 0.2, indicating that these shifts are predominantly independent from the fluctuating tRNA pool (Table S3). This framework is particularly useful for interpreting modifications distributed across multiple tRNA species, like for example the Gm modification. In that case, although tRNA^Ser^(CGA)-1 was downregulated under pH stress at the 24 h timepoint (Fig. S9), the global decrease in Gm (Fig. 3A) exceeded the decline predicted from its carrier abundance (Table S3). Therefore, the *MD_m_* score predicts that the primary cause for the observed change in Gm levels is linked to events that are independent from the decreased tRNA levels.

Furthermore, this quantitative approach indicates that the decrease in the observed thiolation signals is linked tRNA abundance-independent reprogramming events. Although mcm^5^s^2^U could not be directly included in the *MD_m_* analysis in certain stress conditions due to MS detection limits, the *MD_m_* score for its precursor, mcm^5^U, strongly supports that its accumulation is not driven solely by tRNA abundance changes. For example, under heat stress, tRNA^Arg^(UCU) and tRNA^Glu^(UUC), which are carriers of mcm^5^U, showed moderate upregulation in both timepoints (Fig. 4A). However, the *MD_m_* scores for this modification were 0.8200 and 1.0002 at 12 h and 24 h, respectively, indicating that the upregulation of those tRNA carriers is insufficient to account for the pronounced accumulation of mcm^5^U observed by UPLC-MS (Fig. 3A; Table S3). In contrast, mcm^5^s^2^U displayed a slight increase at 12 h under MMS stress (Fig. 3A, B), coinciding with the increased abundance of one of its carriers, tRNA^Lys^(UUU) (Fig. 4A). The corresponding *MD_m_* score for that modification-tRNA molecule pair is 0.0318, indicating that mcm^5^s^2^U in this condition is primarily driven by tRNA abundance. However, the sharp decrease of mcm^5^s^2^U in other tested conditions (heat, pH stress, and paraquat) strongly implies independent reprogramming events rather than tRNA abundance-driven effects. Previous studies have shown that such decreases in thiolation often result from altered activity of the Elongator/Trm9–Trm112 complex and the URM1-dependent thiolation system under heat stress.^16, 35, 41, 43, 44^ Overall, our findings highlight the importance of integrating tRNA abundance changes when interpreting global tRNA epitranscriptomic changes under suboptimal conditions.

### Temporal and condition-specific correlation patterns in tRNA pools

Pairwise Pearson correlation analysis further revealed distinct stress-induced expression relationships (Fig. 4B). At 12 h, most stress conditions showed high similarity in tRNA expression profiles, except for MMS and paraquat, which exhibited weaker correlations with the other stressors. By 24 h, the correlation landscape changed substantially: only heat and suboptimal pH maintained strong similarity, while the other conditions diverged more sharply from one another. When comparing expression profiles across timepoints within each stress, heat and suboptimal pH once again showed the greatest temporal stability, displaying high correlation between 12 h and 24 h (Fig. S6B). In contrast, profiles from MMS and paraquat showed markedly lower cross-timepoint correlation.

## DISCUSSION

In this study, we investigated how diverse environmental and chemical stressors reshape the tRNA epitranscriptome and tRNA expression landscape in *S. cerevisiae*. By combining growth and survival profiling with UPLC-MS and tRNA sequencing, we show that distinct stresses trigger specific, time-dependent tRNA modification signatures. Our *MD_m_* index analysis reveals that these signatures are largely independent from changes in tRNA abundance. Notably, heat stress, paraquat-induced oxidative stress, and pH stress induced a pronounced decrease of wobble uridine thiolation. For heat and pH stress, this decrease was also accompanied by the accumulation of the non-thiolated precursor modification mcm^5^U. This finding is particularly significant given the established role of thiolated tRNA modifications in ensuring translational fidelity through accurate codon recognition while also contributing to cell viability and proteome homeostasis.^3, 18, 35, 41, 43, 55–57^

Our initial phenotypic characterization ensured that the selected stress conditions produce comparable cytotoxic outcomes at selected timepoints. Growth was assessed across a wide range of stress intensities (Fig. 1), and conditions that yielded 60–80% cytotoxicity were chosen for downstream analyses (Fig. 2). This design ensured that cells experienced severe stress that was sufficient to elicit adaptive responses without complete loss of viability. Our late timepoint analyses introduced an additional layer of physiological stress associated with high cell density. Yeast cultures grown to high density or stationary phase experience multiple simultaneous pressures, including nutrient depletion, oxygen limitation, accumulation of toxic metabolites and abnormal proteins, and temperature or osmotic fluctuations.^58, 59^ Moreover, entering the stationary phase further induces defined physiological changes, such as cell cycle arrest, storage carbohydrate accumulation (primarily glycogen and trehalose), and enhanced thermotolerance.^60^ These endogenous stress responses can also affect tRNA modifications: for example, it has been reported that m^5^C levels in tRNA^His^ increase markedly between exponential growth and early to late stationary phases.^61^ Thus, some of the molecular changes observed in our dataset (Fig. 3A) may reflect the combined impact of the stress physiology associated with high-density growth and the applied stressors.

Despite our efforts to produce comparable cytotoxic outcomes, the growth dynamics and survival rate of *S. cerevisiae* varied somewhat. These differences might reflect the extent to which cells can rewire their metabolic and stress-response pathways over time. For instance, growth at pH 7 yielded the lowest viable counts at both early and late timepoints (Fig. 2B, C). As *S. cerevisiae* grows optimally between pH 4 to 6 and actively acidifies its surroundings, exposure to elevated pH constitutes a physiologically relevant stress.^51^ To prevent uncontrolled pH drift, we supplemented the YPD medium with 100 mM KH_2_PO_4_ (Fig. S7). While pH-stat systems have been used in other studies ^51^, supplementing the medium with buffer allowed us to maintain stable pH conditions within our experimental limitations (Fig. S7). On the other hand, acidic pH stress triggered a distinct stress response characterized by a slower growth rate, but faster onset of the exponential growth phase (Fig. 1A, B). It has been shown that acidic stress causes transcriptional changes in genes related to cell wall composition and integrity, oxidation-reduction processes, carbohydrate metabolism, ATP synthesis, and iron uptake.^62^ However, our results showed that long-term acidic stress resulted in pronounced tRNA degradation (Fig. S1), which prevented further modification analysis. Furthermore, the oxidative stressors used in this study elicited distinct physiological responses ^63, 64^—for example, exposure to diamide permitted a partial recovery of the cells, whereas paraquat caused persistent drop in viability (Fig. 2A, B). Diamide is a thiol-specific oxidant that rapidly shifts the glutathione pool toward glutathione disulfide, i.e. the oxidized dimeric form of glutathione, which causes a shift in the cellular redox state.^65^ Consequently, obtaining 60–80 % of reduction of the diamide-stressed cultures was challenging—applying a 2 mM final concentration proved lethal (Fig. 1), while 1.5 mM was insufficient, as the cells recovered quickly and their viable counts was markedly higher than with other stressors (Fig. 2A, B). Indeed, previous studies in mammalian tissues have shown that diamide is rapidly detoxified by the glutathione reductase ^66^, which is consistent with our growth curves and survival data (Fig. 2A, B). In contrast, paraquat engages in a mitochondrial redox cycle that continuously generates superoxides and depletes NADPH.^67^ This sustained oxidative burden explains its higher overall lethality, the rapid decline in survival, and the inability of the cells to recover (Fig. 2A, B). Next, methyl methanesulfonate (MMS), a well-characterized alkylating agent, showed a pronounced cytotoxic effect (Fig. 2A, B). MMS induces DNA damage and suppresses transcription and translation.^68, 69^ In line with its toxicity profile, MMS caused a rapid drop in survival that stabilized without signs of recovery (Fig. 2A, B). Heat stress, however, has been extensively studied in *S. cerevisiae*. Yeast activates a heat shock response (HSR) at temperatures above their optimal growth range (∼30 °C), inducing a broad transcriptional reprogramming that alters cellular physiology to support survival.^70^ Similar to previous observations in *S. cerevisiae* laboratory strains ^16^, our results indicate that temperature over 37 °C significantly impaired growth and viable counts, whereas temperatures over 40 °C led to a complete growth arrest (Fig. 1A).

To cope with this stress-dependent growth inhibition, yeast cells rely on adaptive responses, where tRNA modifications play an essential role, selectively driving translation of codon-biased transcripts essential for survival. ^4, 31–34, 36, 71^ The development of analytical methods that allow precise identification of tRNA modifications has been an active area of research, leading to the advancement of diverse approaches and the continual emergence of new technologies.^72–74^ Nevertheless, liquid chromatography coupled to mass spectrometry of mononucleosides remains the gold standard for tRNA modification analysis, even at the expense of losing sequence context.^75^ Therefore, we profiled the global landscape of tRNA modifications in long-term stressed *S. cerevisiae* cultures using rapid C18-UPLC-MS analysis of enzymatically digested tRNAs, which enabled robust detection of a wide range of modified ribonucleosides.^42^

Our results corroborate prior reports of temperature-sensitive thiolation by demonstrating that prolonged heat stress causes persistent loss of mcm^5^s^2^U (Fig. 3A, Fig. S5A, B). Alings et al. showed that thiolation is impaired at temperatures exceeding 30 °C in laboratory strains S288C and W303 in shorter exposures, which reflects the temperature sensitivity of URM1/Ncs2-Ncs6 thiolation system components rather than defects in the ELP pathway.^16^ They further reported that thiolation recovers following stress relief without evidence for active removal of modified tRNA or selective degradation of thiolated nucleosides.^16^ We confirmed that chronic temperature stress sustains the thiolation impairment, effectively stabilizing in a non-thiolated state over longer timescales (Fig. 3A). As previously reported, we also observe an increase in mcm^5^U (Fig. 3A, B) under prolonged heat exposure.^16^ This might imply that the ELP pathway is active and, without a functional thiolation pathway, it consequently leads to a buildup of mcm^5^U_34_ in tRNAs that would otherwise carry mcm^5^s^2^U_34_.^16^

Furthermore, we observed a partial loss of thiolation in cells exposed to oxidative stress caused by paraquat and pH stress, the latter being partially restored once the cells reach the exponential growth phase after 24 h (Fig. S5A, B). Similarly to temperature stress, we observed that exposure to pH 7 causes a minor increase in mcm^5^U_34_ levels (Fig. 3A). However, we noticed a decline in both mcm^5^s^2^U_34_ and mcm^5^U_34_ levels when the cells were exposed to paraquat. This pattern might reflect a broader stress-associated reduction in the abundance of these wobble uridine modifications, as the global transcriptional and physiological reprogramming induced by oxidative stress.^19, 33^

Modifications at the wobble uridine (U_34_) position of tRNAs have multiple functional roles in eukaryotes. The mcm^5^s^2^U modification, found on tRNA^Lys^(UUU), tRNA^Glu^(UUC), and tRNA^Gln^(UUG), plays a central role in ensuring efficient and accurate translation.^76^ Removal of U_34_ modifications slows translation at cognate codons, induces ribosome pausing, impairs reading frame maintenance and disrupts protein homeostasis, leading to proteotoxic stress and protein aggregation in yeast and metazoans.^3, 55^ Stress-induced remodeling of the tRNA modification landscape extends beyond thiolation, with multiple modifications showing altered levels in both our study and prior work. Previously, it was reported by Alings et al. that a broad reduction in tRNA modification levels is caused by heat stress, particularly affecting wobble-position modifications such as Gm and Cm.^16^ Consistent with this, we also observed a decrease in Cm under the same condition. It is worth noting that comparison of control samples at 12 h and 24 h revealed a general ∼0.5-fold increase in most modifications at 24 h, however Cm showed an approximately two-fold increase (Fig. S3). Consequently, all 24 h stressed samples exhibited significantly reduced abundance of Cm relative to the 24 h control. Given Cm’s low signal intensity, this change should be interpreted cautiously. Next, in response to genotoxic stress induced by MMS, we detected a modest enrichment of several tRNA modifications at 12 h, followed by a global suppression of modification levels at 24 h (Fig. 3A). MMS is also known to methylate nucleobases, including guanine N^7^, adenine N^1^, and cytosine N^3 77^, and short-term MMS exposure has been shown to elevate levels of m^7^G.^32, 33, 78^ However, our dataset did not reveal significant increases in canonical methylated nucleosides such as m^7^G, m^1^A, or m^3^C (Fig. 3A).

To place these stress-induced modification patterns into context, it is necessary to consider the methodological limitations. For instance, we detected a total of 22 modifications, but not all of them are attributable to yeast tRNA. We identified trace amounts of m^6,6^A, a ribosomal RNA modification ^1^, in most of our samples (Fig. S2). Although the tRNA enrichment method used is optimized for rRNA removal, extraction-dependent differences may increase the presence of 5S rRNA.^47^ However, tRNAs are by far the most heavily modified RNA class in the cell ^79^, making any trace contributions from non-tRNA contaminants minimal relative to the abundant tRNA-derived modified nucleosides. Furthermore, we detected m^6^A (Fig. S2), an uncommon modification for tRNA, however its presence is likely due to a known false-positive signal arising from m^1^A undergoing partial Dimroth rearrangement under mildly alkaline conditions.^80^ This artefact might arise during the incubation step used to digest tRNA into mononucleosides, with the combination of elevated temperature and ammonium bicarbonate.^13^

Taking these technical limitations into account, we also evaluated how our results align with previous studies that have reported a higher number of modifications ^16, 33, 42^ compared to our dataset. Our HRMS-based approach provides high mass resolution for confident identification of modification masses, but offers less sensitivity, which can limit the detection of low-abundance species. For example, we did not detect 2’-O-methyluridine, dihydrouridine, or wybutosine in our study. The absence of wybutosine could be explained by its reported instability under acidic conditions ^81, 82^, while the lack of signal for the uridine-based modifications may reflect their poor ionization efficiency ^83, 84^ combined with the limitation of the instrument of choice.

A clearer interpretation of the detected modification changes also requires evaluating whether shifts in tRNA abundance could contribute to the observed modification levels. By implementing the modification deviation (*MD_m_*) score, we developed a quantitative framework that establishes whether the modification change emerges from fluctuations in tRNA abundance (Table S3). Previous works demonstrated that tRNA isoacceptor abundance can be a stress-induced expression response ^53, 54, 85^, supported by showing that tRNA degradation and tRNA gene copy number have a limited contribution ^53^. Consistent with these findings, our observation that most significant modification changes exceed the *MD_m_* > 0.2 threshold provides robust evidence that stress-induced epitranscriptomic reprogramming occurs independently of tRNA abundance fluctuations. This observation extends to the changes in wobble U_34_ modifications—our data shows that the accumulation of mcm^5^U under heat, paraquat, and pH stress cannot be explained by the changes in their carriers (tRNA^Lys^(UUU), tRNA^Glu^(UUC), tRNA^Gln^(UUG) (Table S3). It should be noted that the *MD_m_* index currently utilizes a subset of 47 cytoplasmic and mitochondrial isodecoders with annotated modification sites in tModBase ^86^, which account to 63 % of the total isodecoders found in yeast. The current *MD_m_* index is therefore relative to the included tRNA species rather than the absolute total pool, which may oversimplify the interpretation of the relationship tRNA modifications and their carriers under long-term environmental pressure.

Despite that tRNA abundance shifts were generally insufficient to explain global modification shifts, we observed stress-related changes in tRNA abundance in some isoacceptors. Unlike previous short-term stress studies where authors used diauxic, heat, osmotic and oxidative stress ^54^, we did not observe a consistent decrease in tRNAs encoding glutathione-related amino acids glutamic acid, cysteine and glycine (Fig. 4A). This possibly reflects the difference in stress duration, as several tRNA isoacceptor trends in our long-term stress dataset show opposite regulation patterns (Fig. 4A). For example, serine and leucine isoacceptors, both consistently upregulated in Pang et al. and Torrent et al. under acute stress, were markedly downregulated in our data.^53, 54^ In contrast, arginine tRNAs levels were uniformly elevated in our long-term stress data, but in other studies displayed mixed responses (Fig. 4A).^53, 54^

Ultimately, by providing a framework to decouple the influence of tRNA abundance from global modification signals, our study establishes that long-term environmental stress triggers targeted reprogramming events of the yeast tRNA epitranscriptome independent from tRNA abundance as a primary adaptive strategy.

## CONCLUSIONS

This study provides the first comprehensive atlas of global tRNA modification and tRNA isoacceptor abundance changes in *S. cerevisiae* under sustained environmental stress. Here, we demonstrate that long-term stress triggers a global, stress-specific, and time-dependent reprogramming of the tRNA epitranscriptome. By implementing the *MD_m_* score to integrate mass spectrometry-based and tRNA-seq data, we show these shifts are mostly independent from fluctuations in tRNA abundance. Furthermore, a central discovery of this work is the novel identification of complete or partial loss of wobble uridine thiolation (mcm^5^s^2^U_34_) under oxidative (paraquat) and pH (pH 7) stress, which previously has been reported only for heat stress. Moreover, this decrease in thiolation signal is accompanied by the specific accumulation of the non-thiolated mcm^5^U_34_ precursor modification. These findings further clarify the mechanism behind these changes, showing that the observed tRNA modification signatures are independent from changes in the expression levels of the tRNA isoacceptors, confirmed by the *MD_m_* scores. By addressing the critical gap in knowledge regarding prolonged stress exposure, this study enhances our understanding of translational control and provides a foundation for engineering stress-resilient yeast strains to optimize protein yields and robustness in industrial biotechnology.

## ASSOCIATED CONTENT

### Supporting Information

The following files are available free of charge.

Supplemental figures (S1-S9): tRNA isolation on urea-polyacrylamide gel (S1), additional plots related to UPLC-MS analysis (S2-S4), northern blot confirming thiolation loss (S5), pairwise correlation of changes in tRNA modifications between two time points (S6), buffer adjustment for basic and alkaline pH (S7), additional plots related to tRNA-seq analysis (S8, S9); supplemental tables (S1-S4): cytotoxicity screening of stressor exposure in *S. cerevisiae* (S1), relative (fold) change in nucleoside modification levels (S2), *MD_m_* scores (S3), DNA oligonucleotide probes used for northern blotting (S4), references (PDF)

*MD_m_* calculation, 12 h (Excel)

*MD_m_* calculation, 24 h (Excel)

## Data Availability Statement

The data that support this study are available from the corresponding author upon reasonable request. All tRNA-seq raw files generated during this study are available for download on the NCBI Sequence Read Archive (SRA) database under BioProject accession number PRJNA1443207.

## AUTHOR INFORMATION

Present Addresses: †If an author’s address is different than the one given in the affiliation line, this information may be included here. Matea Radešić – RNAcious Laboratory, Department of Molecular and Integrative Biosciences, Faculty of Biological and Environmental Sciences, University of Helsinki, 00014 Helsinki, Finland; Doctoral Programme in Integrative Life Science, University of Helsinki Doctoral School, University of Helsinki, 00014 Helsinki, Finland;

Jenni K. Pedor – RNAcious Laboratory, Department of Molecular and Integrative Biosciences, Faculty of Biological and Environmental Sciences, University of Helsinki, 00014 Helsinki, Finland; Doctoral Programme in Integrative Life Science, University of Helsinki Doctoral School, University of Helsinki, 00014 Helsinki, Finland;

M. Suleman Qasim – RNAcious Laboratory, Department of Molecular and Integrative Biosciences, Faculty of Biological and Environmental Sciences, University of Helsinki, 00014 Helsinki, Finland; Doctoral Programme in Microbiology and Biotechnology, University of Helsinki Doctoral School, University of Helsinki, 00014 Helsinki, Finland;

Anna-Emilia Rajaveräjä – RNAcious Laboratory, Department of Molecular and Integrative Biosciences, Faculty of Biological and Environmental Sciences, University of Helsinki, 00014 Helsinki, Finland;

Nina Sipari – School of Pharmacy, Faculty of Health Sciences, University of Eastern Finland, 70211 Kuopio, Finland;

## Author Contributions

M.R. contributed to conceptualization, methodology, formal analysis, investigation, data curation, writing of the original draft, writing (review and editing), and visualization. J.K.P. contributed to methodology, investigation, data curation, and writing (review and editing).

M.S.Q. contributed to methodology, formal analysis, data curation, and writing (review and editing). A-E.R. contributed to investigation, data curation, and writing (review and editing).

N.H.S. contributed to methodology, investigation, and writing (review and editing). L.P.S. contributed to conceptualization, resources, supervision, project administration, funding acquisition, and writing (review and editing). All authors have read and approved the final version of the manuscript.

## Funding Sources

This research was funded by the Research Council of Finland (grant #354906; to L.P.S.) and the Novo Nordisk Foundation (grant #NNF19OC0054454; to L.P.S.). M.R. and J.K.P. are fellows of the Doctoral Programme in Molecular and Cellular Systems of Life, University of Helsinki, and M.S.Q. is a fellow of the Doctoral Programme in Microbiology and Biotechnology, University of Helsinki.

## ACKNOWLEDGMENTS

The authors thank Salla Kalaniemi and Sari Korhonen for their valuable technical assistance. The authors wish to acknowledge the Instruct-HiLIFE Biocomplex Unit at the University of Helsinki, a member of Instruct-ERIC Centre Finland, FINStruct, and Biocenter Finland for centrifugation services, and the Next Generation Sequencing Facility at Vienna BioCenter Core Facilities (VBCF), member of the Vienna BioCenter (VBC), Austria, for NGS services. CSC – IT Center for Science Ltd., Finland, is acknowledged for computational resources. The authors are grateful to Miikka Olin, Department of Food and Nutrition, Faculty of Agriculture and Forestry, University of Helsinki for the use of their Synapt G2 Si mass spectrometer. Pavlina Gregorova is thanked for the design of the DNA oligonucleotide probes used in the Northern Blot experiment. Lastly, we thank all members of the RNAcious laboratory for their insightful feedback and supportive discussions. The graphical abstract was created with Biorender. The authors acknowledge the use of Perplexity Pro and M365 Copilot for language editing and proofreading during the preparation of this manuscript.

## ABBREVIATIONS

C: cytidine
Ψ: pseudouridine
m^3^C: 3-methylcytidine
m^1^A: 1-methyladenosine
m^5^C: 5-methylcytidine
U: uridine
m^7^G: 7-methylguanosine
Cm: 2’-O-methylcytidine
A: adenosine
G: guanosine
I: inosine
m^5^U: 5-methyluridine
Am: 2’-O-methyladenosine
m^1^G: 1-methylguanosine
Gm: 2’-O-methylguanosine
m^2^G: 2-methylguanosine
m^6^A: 6-methyladenosine
m^1,3^Ψ: 1,3-dimethylpseudouridine
mcm^5^U: 5-methoxycarbonylmethyluridine
m^2,2^G: N2,N2-dimethylguanosine
mcm^5^s^2^U: 5-methoxycarbonylmethyl-2-thiouridine
m^1^I: 1-methylinosine
ncm^5^U: 5-carbamoylmethyluridine
t^6^A: N6-threonylcarbamoyladenosine
i^6^A: N6-isopentenyladenosine.

## Notes

### Competing Interest Statement

The authors have declared no competing interest.

### Summary of Updates

The abstract has been rewritten and the manuscript text updated throughout. Minor edits have been performed to the figures and the supplementary information document has been thoroughly revised.

