## Supplementary Information for "Atlas of stress-induced changes in yeast transfer RNA modification levels"

Table of contents

Figure S1. tRNA isolation on urea-polyacrylamide gel across conditions ........ S2

Figure S2. Heatmap of nucleoside ions detected in stressed *S. cerevisiae* cultures by UPLC-MS. ........ S3

Figure S3. Fold changes of modified nucleosides between control samples at 24 h and 12 h quantified by UPLC-MS. ........ S4

Figure S4. Volcano plots show the distribution of tRNA modifications in response to stressors for 12 h (upper panels) and 24 h (lower panels) ........ S5

Figure S5. Northern blot validation of condition-specific thiolation loss. ........ S6

Figure S6. Bar plots showing pairwise correlation of changes in tRNA modifications between two time points (12 h and 24 h) in stressors applied to *S. cerevisiae* cultures ........ S7

Figure S7. Adjustment of buffer concentration for alkaline pH YPD. ........ S8

Figure S8. Differential expression analysis of cytosolic *S. cerevisiae* tRNA isoacceptors transcripts (counts) in control samples at 12 h and 24 h. ........ S9

Figure S9. Changes in tRNA isodecoders expression under selected stress conditions ........ S10

Table S1. Cytotoxicity screening of stressor exposure in *S. cerevisiae* ........ S11

Table S2. $Log_2$ fold change in nucleoside modification levels compared to control, marked by significance[a] with standard deviation (+/-)[b]. ........ S12

Table S3. Quantitative assessment of abundance-driven tRNA modification regulation by modification deviation ($MDm$) scores[a,b]. ........ S14

Table S4. DNA oligonucleotide probes used for northern blotting. ........ S15

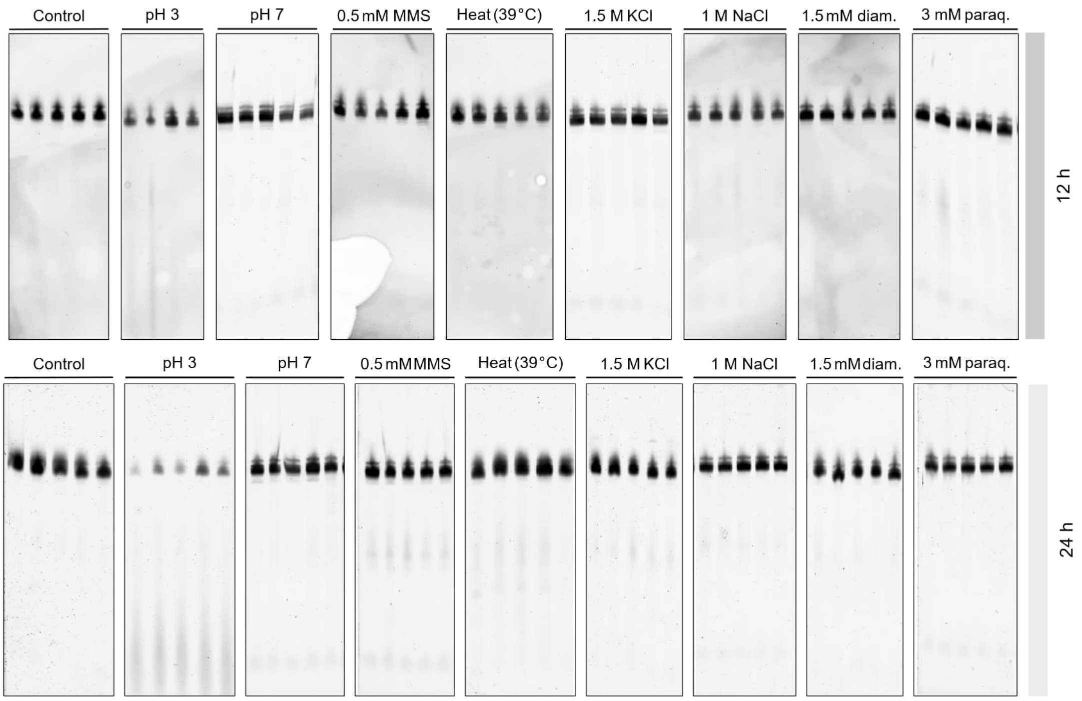


Figure S1. tRNA isolation on urea-polyacrylamide gel across conditions. Representative 10 % urea-polyacrylamide gel showing tRNA isolated from *S. cerevisiae* grown under stress conditions after 12 hours (top panel) and 24 hours (bottom panel) of exposure. Each panel contains lanes corresponding to the following conditions, as labeled: control (untreated cultures), pH 3, pH 7, 0.5 mM MMS, heat (39 °C), 1.5 M KCl, 1 M NaCl, 1.5 mM diamide, and 3 mM paraquat. Consistent degradation is visible in tRNA samples isolated from cultures grown in pH 3, which were omitted for further analyses. Data is shown for 5 biological replicates, except for pH 3, 12 h, where 4 biological replicates were used.

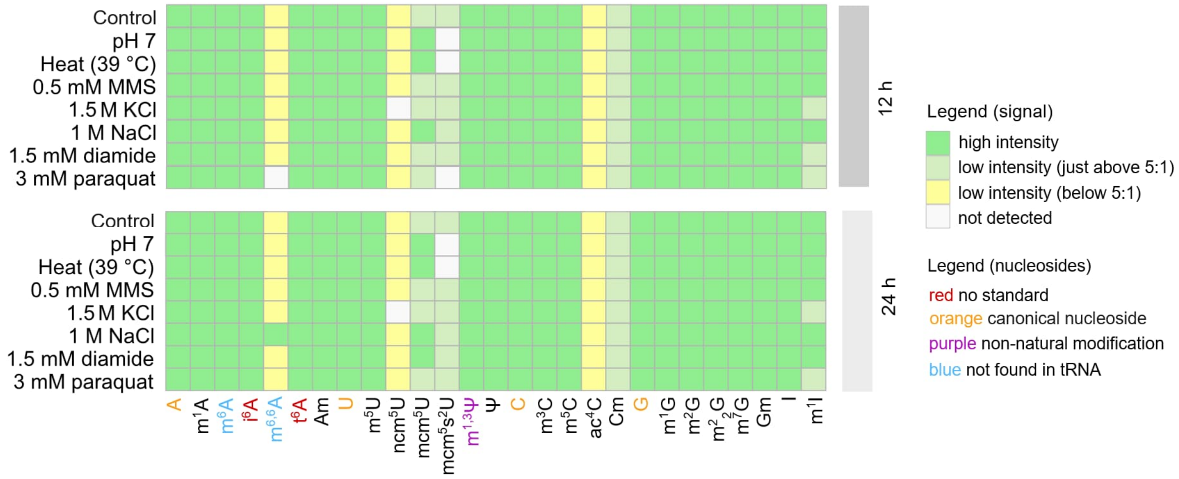


Figure S2. Heatmap of nucleoside ions detected in stressed *S. cerevisiae* cultures by UPLC-MS. The heatmap visualizes the signal robustness of ions identified as nucleosides across different conditions. Each column represents a distinct tRNA modification, and each row corresponds to a specific stress condition. Ion signals are categorized according to their signal-to-noise (S/N) ratio: green – robust, high-intensity signals (≫5:1) suitable for comparative analysis, light green – detectable signals with low-abundance (just above 5:1), yellow – low-intensity signals that fell below the established threshold for characterization; and grey – ions that were not detected. Text color highlights the annotation category for each modification: red – modifications for which analytical standards were unavailable (identity inferred by mass); orange – canonical nucleosides; purple – non-natural internal standard used for normalization; and blue – modifications not typically found in tRNA, which may result from contamination or analytical artifacts.

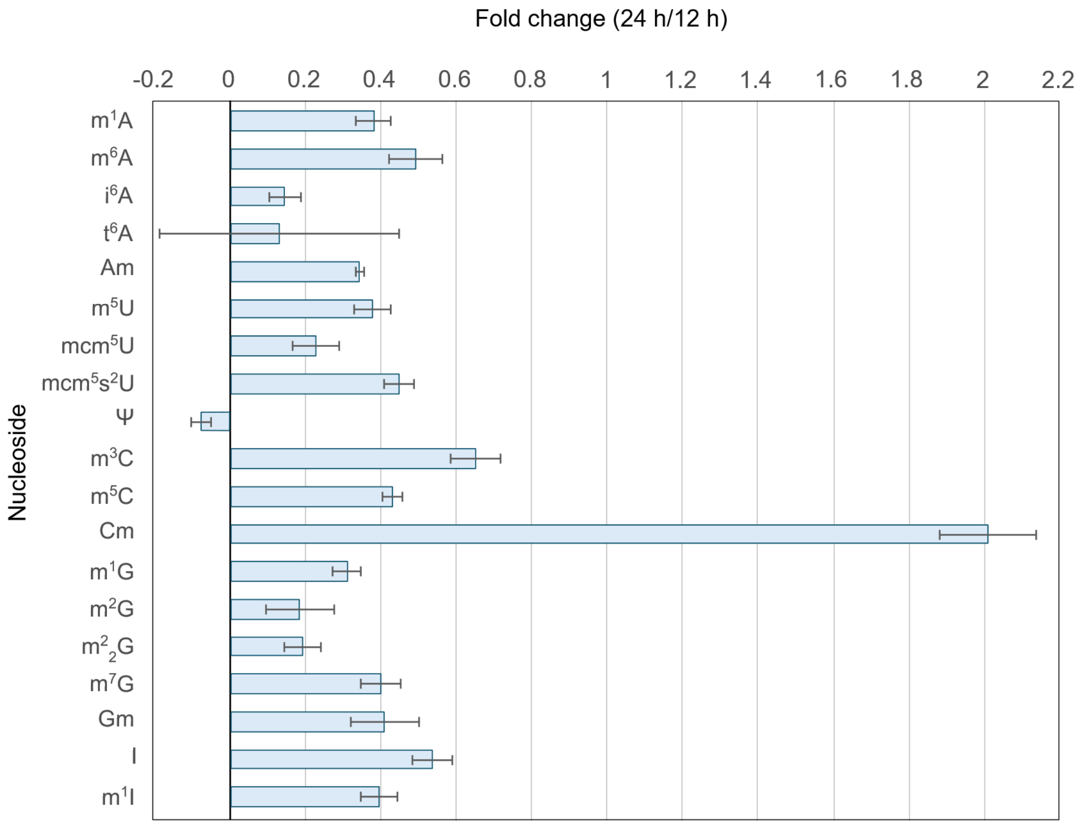


Figure S3. Fold changes of modified nucleosides between control samples at 24 h and 12 h quantified by UPLC-MS. The bar graph shows centered fold changes (i.e., by subtracting 1) of individual modified nucleosides (24 h relative to 12 h) in control samples. UPLC-MS peak height signals were normalized to the internal standard 1,3-dimethylpseudouridine. Each bar represents the mean centered fold change for a given nucleoside, with error bars indicating the standard deviation across five biological replicates. Most nucleosides show only small time-dependent shifts, whereas Cm displays a notably larger increase.

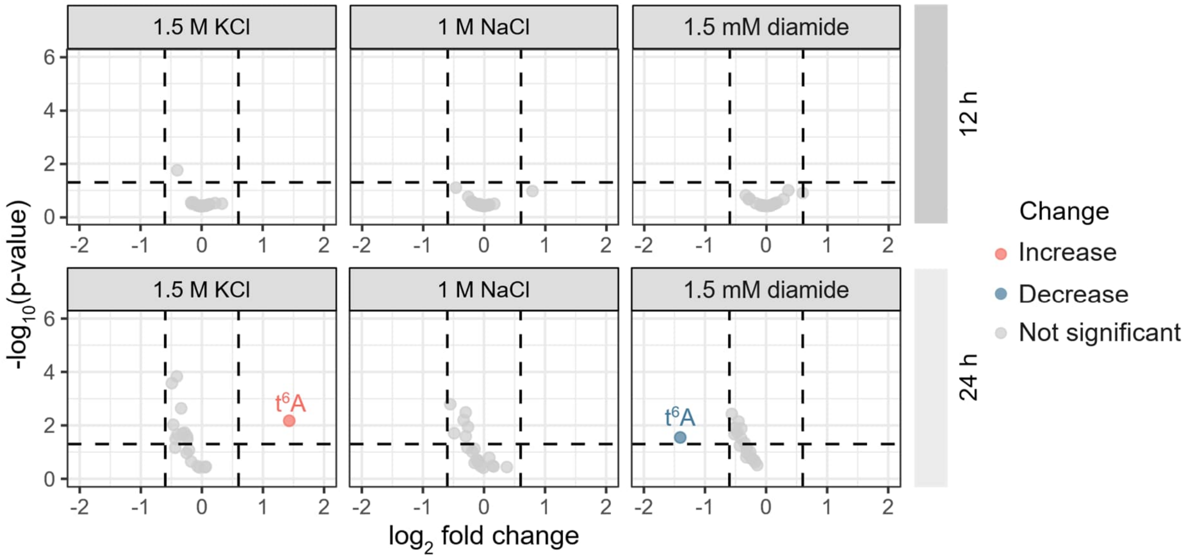


Figure S4. Volcano plots show the distribution of tRNA modifications in response to stressors for 12 h (upper panels) and 24 h (lower panels). Each point represents a tRNA modification, plotted by $\log_2$ fold change (x-axis) and statistical significance as $-\log_{10}$ of p-value (y-axis). P-values were calculated using an unpaired two-samples *t*-test. Points meeting significance criteria ($|\log_2 \text{fold}| > 0.6$ and $p < 0.05$) are colored in coral red – increased levels compared to the control, and blue – decreased levels compared to the control. Non-significant points are shown in grey. The volcano plots display significant changes present only for modification $t^6A$, without a standard for mass spectrometry analysis. Dashed vertical lines mark fold change thresholds (± 0.6), and the dashed horizontal line marks the significance threshold ($p = 0.05$). Data are derived from UPLC-MS analysis across 5 biological replicates, except for 1.5 M NaCl, 24 h (n = 3) and 1 M KCl, 12 h (n = 4).

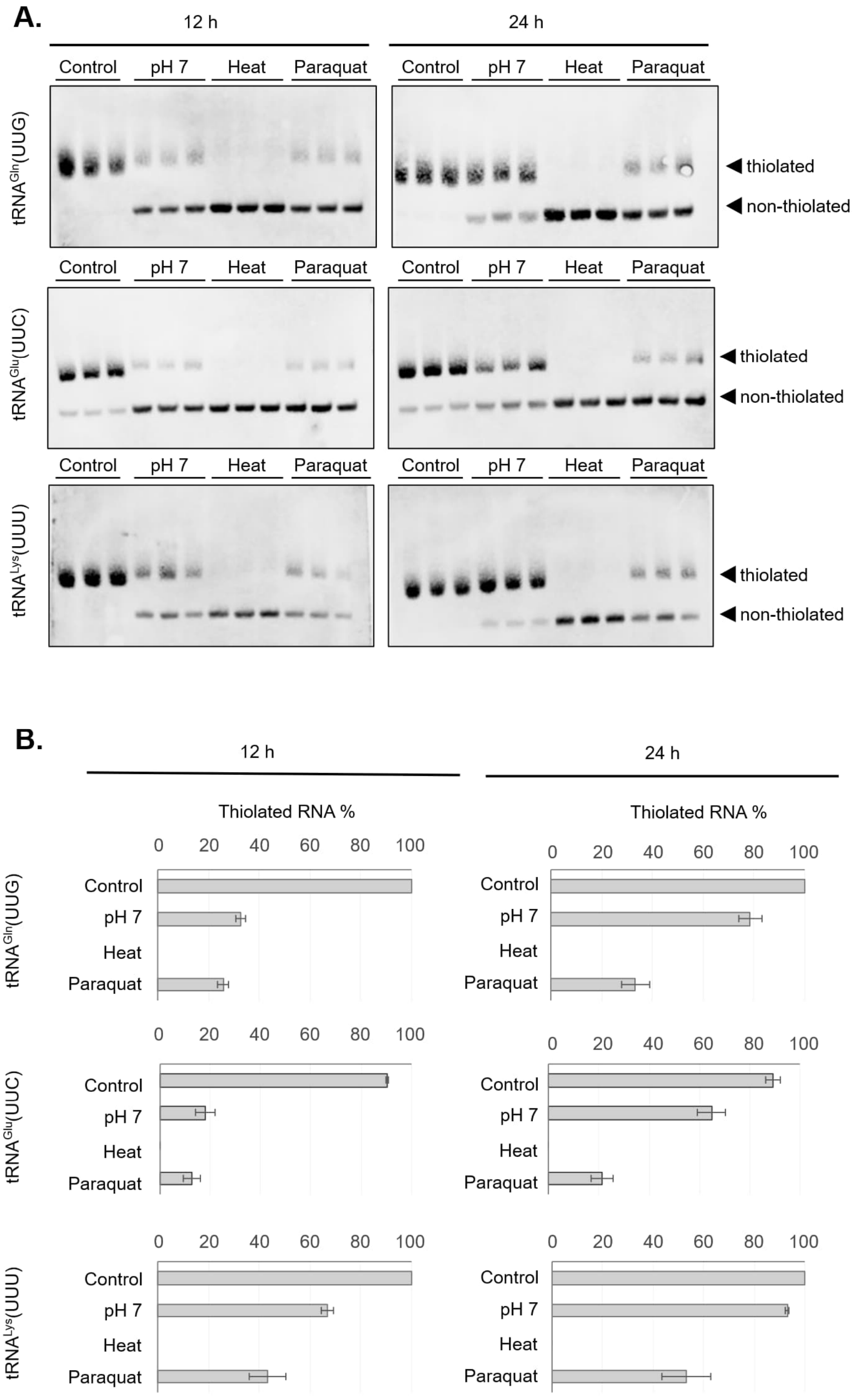


Figure S5. Northern blot validation of condition-specific thiolation loss. (A) Northern blots of bulk tRNA isolated from *S. cerevisiae* cultures after different stress exposure for 12 h (left) or 24 h (right) following APM-affinity gel electrophoresis. The tested biotinylated DNA probes are marked on the left. On the right, the horizontal arrows mark thiolated and non-thiolated tRNA species. (B) Quantification of thiolation levels by image densitometry (n = 3) performed in ImageLab (v. 6.0).

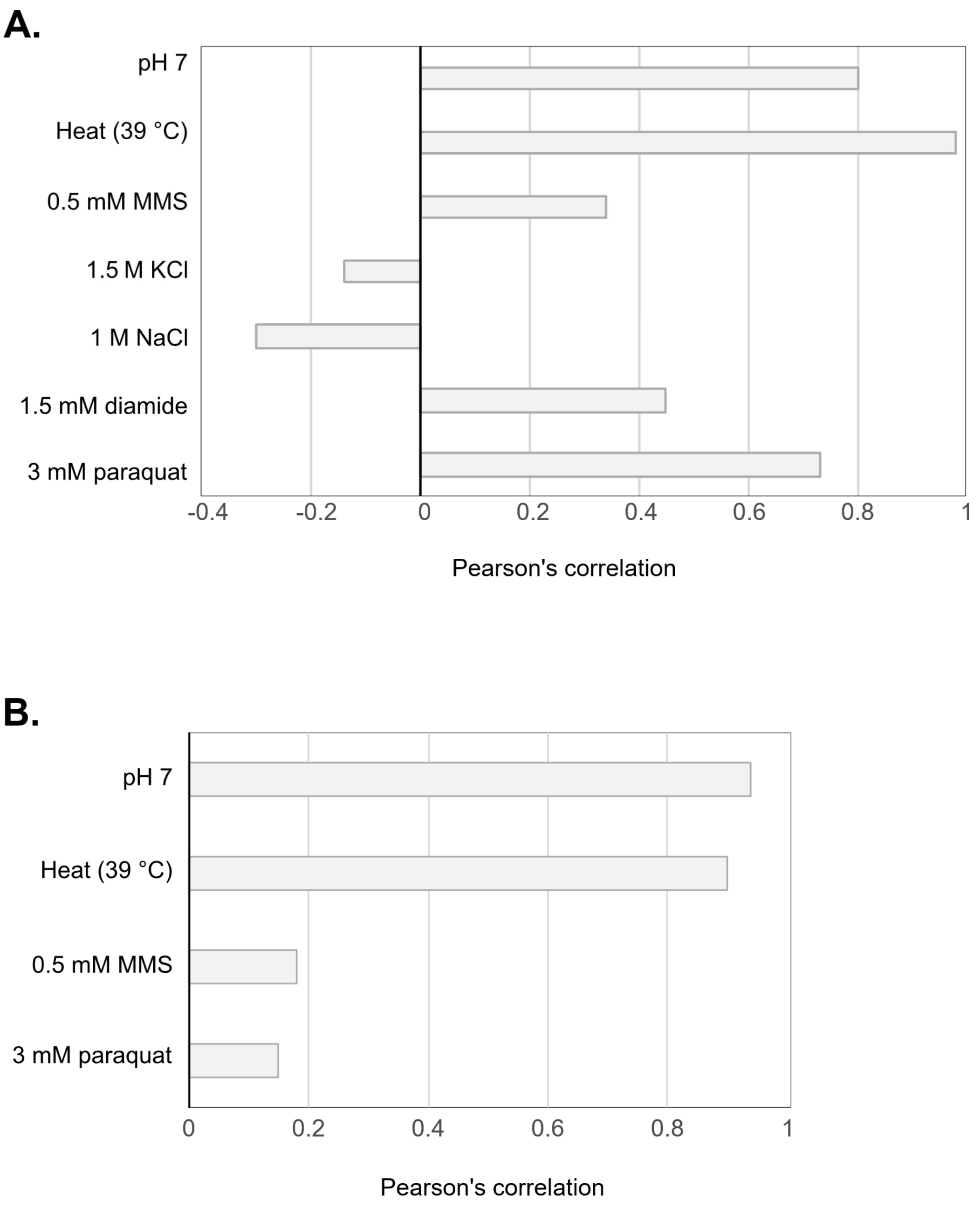


Figure S6. Bar plots showing pairwise correlation of changes in tRNA modifications between two time points (12 h and 24 h) in stressors applied to *S. cerevisiae* cultures. (A) Pairwise correlations of stress-induced changes in tRNA modifications measured at 12 h and 24 h. Each bar represents the Pearson correlation coefficient (*r*) for a given stress condition, calculated using the dataset shown in Fig. 3A. (B) Pairwise correlations of stress-induced changes in tRNA isoacceptors measured at 12 h and 24 h. Each bar represents the Pearson correlation coefficient (*r*) for the corresponding stress condition, calculated from the tRNA-seq data shown in Fig. 4A.

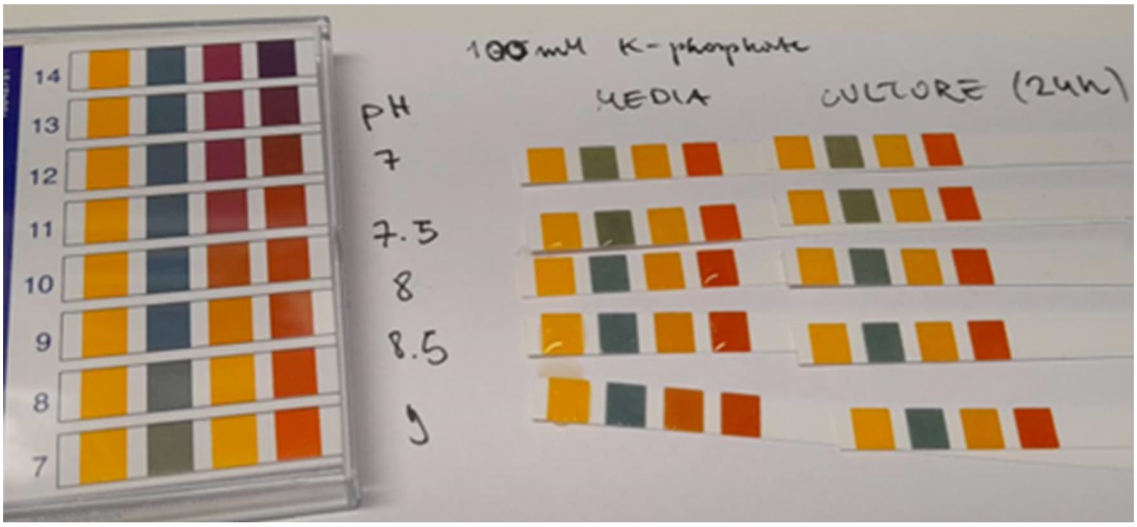


Figure S7. Adjustment of buffer concentration for alkaline pH YPD. Litmus paper pH scale (left), YPD adjusted to pH 7–9, supplemented with 100 mM $KH_2PO_4$ soaked papers (middle), and *S. cerevisiae* cultures grown in YPD adjusted to pH 7–9, supplemented with 100 mM $KH_2PO_4$ after 24 h (right).

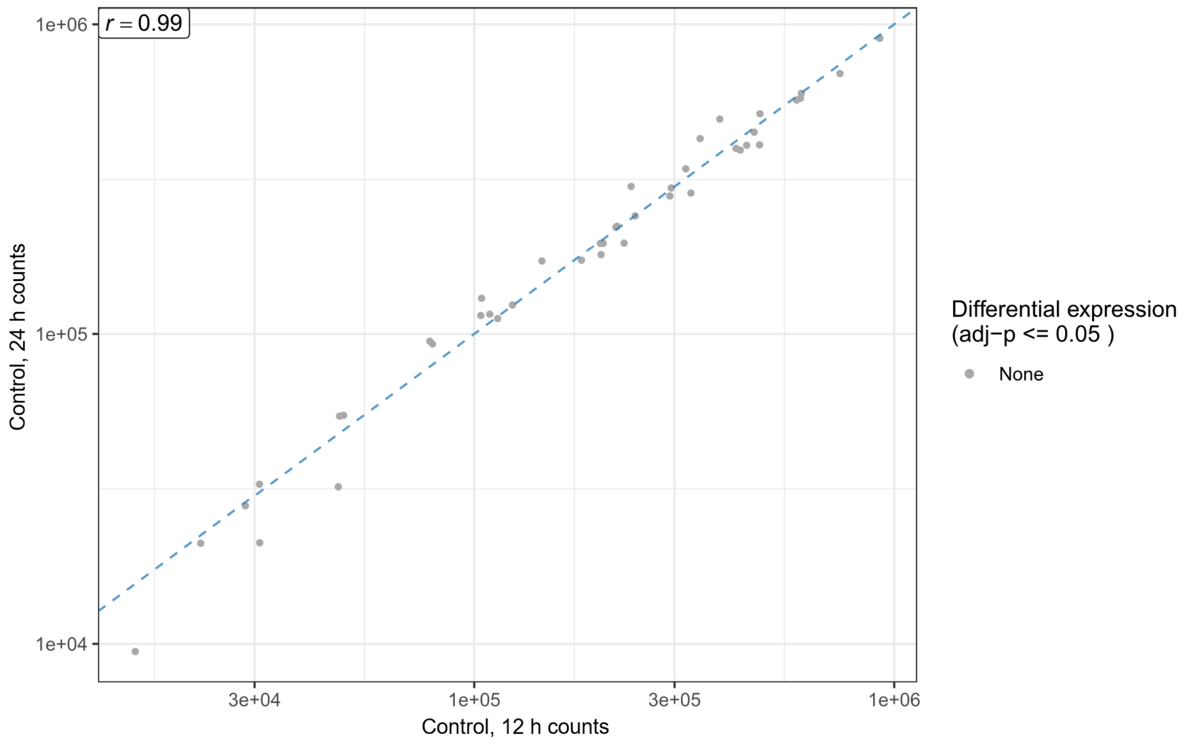


Figure S8. Differential expression analysis of cytosolic *S. cerevisiae* tRNA isoacceptors transcripts (counts) in control samples at 12 h and 24 h. Axes represent log-transformed normalized read counts from DESeq2.

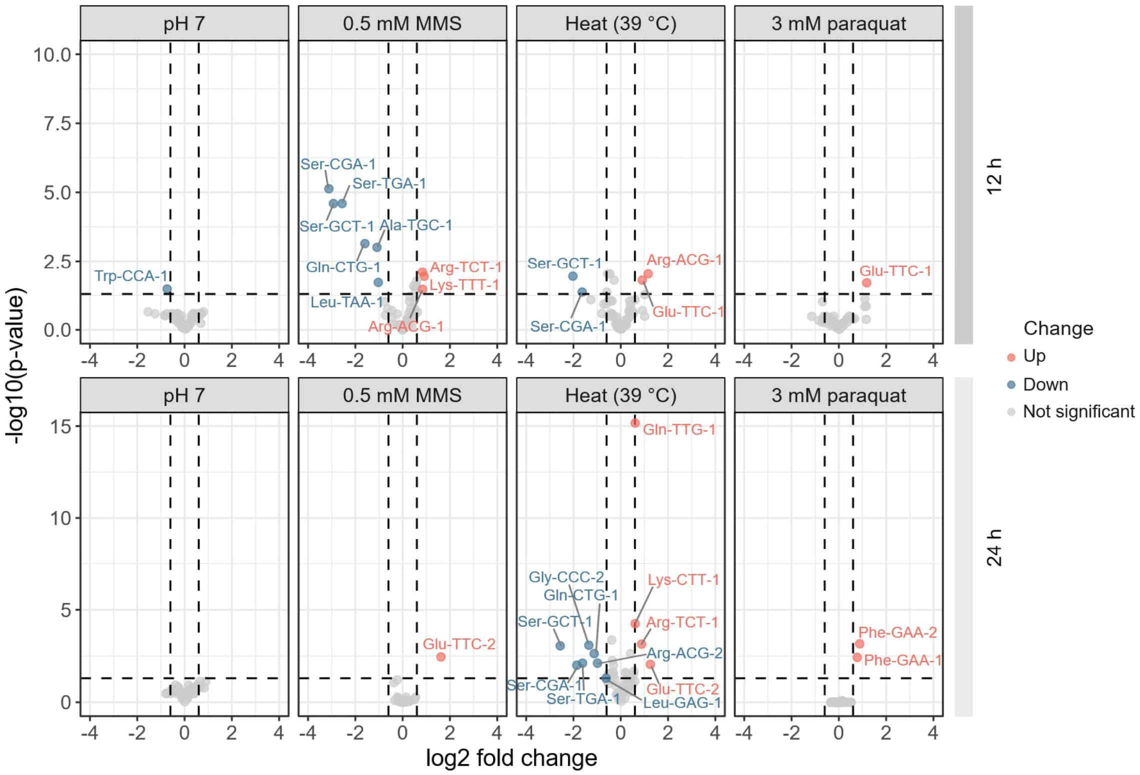


Figure S9. Changes in tRNA isodecoders expression under selected stress conditions. Volcano plots show the distribution of tRNA isodecoders in response to each stress exposure for 12 h and 24 h. Each point represents a tRNA isodecoder, plotted by $log_2$ fold change (x-axis) and statistical significance as -$log_{10}$ of p-value (y-axis). Points meeting significance criteria ($|log_2 fold| > 0.6$ and $p < 0.05$) are colored in coral red (upregulation) or blue (downregulation). Non-significant points are shown in grey. Dashed vertical lines mark fold change thresholds (± ~1.5), and the dashed horizontal line marks the significance threshold ($p = 0.05$). Data are derived from tRNA-seq analysis across independent biological replicates ($n = 5$), except for samples where $n = 4$ (control at 12 h) and $n = 3$ (paraquat at 12 h).

Table S1. Cytotoxicity screening of stressor exposure in *S. cerevisiae*.

| Stressor | Stressor dose at screening phase |
|---|---|
| Heat | 37, 39, 40, 42 °C |
| Osmotic | 1, 1.5, 2 M NaCl/KCl |
| Genotoxic | 1, 2, 3.5, 5, 6 mM MMS |
| Acidic | pH 3, 3.5, 4 ($H_3PO_4$) |
| Neutral and alkaline | pH 7, 7.5, 8, 8.5, 6 (KOH) + 100 mM $KH_2PO_4$ |
| Oxidative | 1, 2, 3.5, 5, 6 mM paraquat; 1, 2, 3.5, 5 mM diamide |

Table S2. $Log_2$ fold change in nucleoside modification levels compared to control, marked by significance[a] with standard deviation (+/-)[b].

| Modification | pH 7 | +/- | 0.5 mM MMS | +/- | Heat 39 °C | +/- | 1.5 M KCl | +/- | 1 M NaCl | +/- | 1.5 mM diamide | +/- | 3 mM paraquat | +/- |
|---|---|---|---|---|---|---|---|---|---|---|---|---|---|---|
| 12 h | | | | | | | | | | | | | | |
| m$^1$A | 0.0307 | 0.0833 | 0.3431 | 0.0671 | -0.1561 | 0.0869 | -0.0135 | 0.0830 | -0.2133 | 0.0869 | -0.0451 | 0.0298 | -0.0795 | 0.0824 |
| m$^6$A | 0.1004 | 0.1184 | 0.4687 | 0.0824 | -0.1746 | 0.0944 | -0.0461 | 0.0923 | -0.0225 | 0.1202 | 0.0670 | 0.0633 | -0.1515 | 0.1683 |
| i$^6$A | -0.0792 | 0.1554 | 0.5005 | 0.5372 | -0.3143 | 0.1382 | -0.0123 | 0.1702 | -0.1185 | 0.1112 | -0.1764 | 0.0587 | -0.0629 | 0.0693 |
| t$^6$A | 0.5036 | 0.3296 | 0.2896 | 0.5372 | 0.2380 | 0.4699 | 0.3305 | 0.4407 | 0.7876 | 0.3051 | 0.5941 | 0.2677 | 0.9899 | 0.2223 |
| Am | 0.1065 | 0.0134 | 0.3122 | 0.0232 | 0.0432 | 0.0154 | 0.0736 | 0.0288 | -0.0809 | 0.0168 | 0.1397 | 0.0327 | 0.1499 | 0.0143 |
| m$^5$U | 0.1648 | 0.0695 | 0.4047 | 0.0621 | -0.2195 | 0.0739 | 0.0718 | 0.0933 | -0.0413 | 0.0252 | -0.0463 | 0.0537 | 0.0857 | 0.1183 |
| mcm$^5$U | 0.5664 | 0.0516 | 0.4710 | 0.0656 | 1.2290 | 0.0468 | -0.1362 | 0.0460 | 0.0770 | 0.0356 | 0.0784 | 0.0902 | -0.8104 | 0.0532 |
| mcm$^5$s$^2$U | -0.3691 | 0.0559 | 0.7315 | 0.0556 | n/d | - | -0.3975 | 0.0247 | 0.0314 | 0.0141 | -0.0936 | 0.1093 | n/d | - |
| Ψ | 0.1906 | 0.0708 | 0.5429 | 0.0495 | -0.0494 | 0.0553 | -0.1622 | 0.1097 | -0.4654 | 0.0857 | 0.0201 | 0.0811 | -0.2576 | 0.1795 |
| m$^3$C | 0.2639 | 0.0853 | 0.5067 | 0.0680 | -0.0581 | 0.0916 | 0.0305 | 0.1046 | -0.1991 | 0.0878 | 0.3582 | 0.0754 | 0.2845 | 0.0261 |
| m$^5$C | 0.1067 | 0.0804 | 0.3213 | 0.0520 | -0.0294 | 0.0559 | -0.1695 | 0.0763 | -0.2637 | 0.0572 | -0.0362 | 0.0400 | -0.1263 | 0.0976 |
| Cm | 0.0337 | 0.1254 | 0.3115 | 0.0825 | -0.1130 | 0.0608 | 0.0685 | 0.1128 | -0.1539 | 0.0915 | -0.2920 | 0.1201 | 0.5387 | 0.1152 |
| m$^1$G | 0.4289 | 0.0697 | 0.5554 | 0.0569 | 0.0392 | 0.0337 | 0.1277 | 0.0898 | 0.0060 | 0.0316 | 0.1252 | 0.0727 | 0.2827 | 0.1068 |
| m$^2$G | -0.1727 | 0.1171 | 0.2228 | 0.0668 | -0.1775 | 0.1159 | -0.0911 | 0.0830 | -0.1588 | 0.0869 | -0.2767 | 0.0630 | -0.3016 | 0.0961 |
| m$^2_2$G | -0.0533 | 0.0644 | 0.2519 | 0.0581 | -0.1085 | 0.0684 | 0.0477 | 0.1158 | -0.0444 | 0.0164 | -0.0963 | 0.0787 | -0.2056 | 0.0915 |
| m$^7$G | 0.0347 | 0.0958 | 0.4343 | 0.0822 | -0.0651 | 0.1056 | -0.0051 | 0.1072 | -0.1948 | 0.0743 | 0.0856 | 0.0598 | -0.0708 | 0.0879 |
| Gm | -0.3068 | 0.1250 | 0.0713 | 0.0975 | -0.2330 | 0.1068 | -0.0454 | 0.1146 | -0.0726 | 0.0652 | -0.3407 | 0.0256 | -0.3449 | 0.0532 |
| I | 0.1330 | 0.0907 | 0.5178 | 0.0937 | 0.0355 | 0.1210 | 0.1215 | 0.0955 | 0.0745 | 0.0562 | 0.1726 | 0.0755 | 0.0453 | 0.1062 |
| m$^1$I | 0.5403 | 0.0665 | 0.7858 | 0.1239 | 0.3093 | 0.1945 | 0.2192 | 0.1537 | 0.1618 | 0.1057 | 0.2807 | 0.1187 | 0.3229 | 0.1245 |
| 24 h | | | | | | | | | | | | | | |
| m$^1$A | -0.6245 | 0.0289 | -0.4821 | 0.0282 | -0.3753 | 0.0206 | -0.2457 | 0.0137 | -0.1917 | 0.0231 | -0.3137 | 0.0412 | -0.3152 | 0.0215 |
| m$^6$A | -0.6215 | 0.0311 | -0.3086 | 0.0465 | -0.7169 | 0.0427 | -0.4690 | 0.0380 | -0.0581 | 0.0195 | -0.5147 | 0.0495 | -0.1385 | 0.0302 |
| i$^6$A | -0.3794 | 0.0356 | -0.2552 | 0.0310 | -0.6072 | 0.0286 | -0.3399 | 0.0110 | 0.0865 | 0.0054 | -0.4510 | 0.0296 | -0.7601 | 0.0440 |
| t$^6$A | 0.1113 | 0.3954 | -0.8623 | 0.2702 | 0.2441 | 0.2242 | 1.4300 | 0.2597 | 0.1575 | 0.1850 | -1.4049 | 0.4923 | 0.4272 | 0.2553 |
| Am | -0.4396 | 0.0529 | -0.8070 | 0.0328 | -0.3912 | 0.0208 | 0.0630 | 0.0345 | -0.2983 | 0.0231 | -0.2353 | 0.0526 | -0.4517 | 0.0217 |
| m$^5$U | -0.3583 | 0.0222 | -0.3787 | 0.0225 | -0.5264 | 0.0208 | -0.3147 | 0.0215 | -0.1195 | 0.0157 | -0.4091 | 0.0460 | -0.2417 | 0.0273 |
| mcm$^5$U | 0.1384 | 0.1179 | -0.1937 | 0.0264 | 1.0267 | 0.0390 | -0.4309 | 0.0331 | -0.1535 | 0.0433 | -0.3231 | 0.1011 | -0.8062 | 0.0807 |
| mcm$^5$s$^2$U | -0.9888 | 0.1030 | -0.9282 | 0.0247 | n/d | - | 0.0503 | 0.0557 | -0.4886 | 0.0546 | -0.4293 | 0.0718 | n/d | - |
| Ψ | -0.2604 | 0.0570 | -0.2976 | 0.0106 | -0.3350 | 0.0237 | -0.4372 | 0.0625 | 0.1434 | 0.0627 | -0.1802 | 0.0474 | -0.3422 | 0.0311 |
| m$^3$C | -0.5838 | 0.0454 | -0.3905 | 0.0284 | -0.3142 | 0.0269 | -0.2497 | 0.0395 | -0.5518 | 0.0254 | -0.1982 | 0.0480 | -0.4304 | 0.0253 |
| m$^5$C | -0.5419 | 0.0147 | -0.6560 | 0.0114 | -0.4758 | 0.0181 | -0.4105 | 0.0077 | -0.3333 | 0.0145 | -0.5610 | 0.0277 | -0.4877 | 0.0121 |
| Cm | -1.3869 | 0.0340 | -1.0859 | 0.0459 | -0.6034 | 0.0241 | -0.3961 | 0.0380 | 0.3741 | 0.2894 | -0.4066 | 0.0325 | -0.6628 | 0.0365 |
| m$^1$G | -0.0730 | 0.0319 | -0.3521 | 0.0405 | -0.3150 | 0.0179 | -0.2123 | 0.0251 | -0.0878 | 0.0061 | -0.2588 | 0.0380 | -0.0282 | 0.0265 |
| m$^2$G | -0.5344 | 0.0481 | -0.4562 | 0.0594 | -0.3117 | 0.0340 | -0.0371 | 0.0695 | -0.0109 | 0.0232 | -0.3150 | 0.0592 | -0.3217 | 0.0613 |
| m$^2_2$G | -0.3640 | 0.0501 | -0.6381 | 0.0404 | -0.3721 | 0.0131 | -0.0718 | 0.0379 | -0.1020 | 0.0062 | -0.2632 | 0.0554 | -0.2781 | 0.0315 |
| m$^7$G | -0.5152 | 0.0247 | -0.4637 | 0.0259 | -0.5908 | 0.0152 | -0.2813 | 0.0158 | -0.1566 | 0.0032 | -0.3519 | 0.0405 | -0.3603 | 0.0296 |
| Gm | -0.8220 | 0.0255 | -0.7975 | 0.0527 | -0.4921 | 0.0223 | -0.1759 | 0.0522 | -0.2708 | 0.0209 | -0.4737 | 0.0461 | -0.4199 | 0.0444 |
| I | -0.5852 | 0.0238 | -0.5251 | 0.0104 | -0.5695 | 0.0214 | -0.4921 | 0.0101 | -0.2994 | 0.0071 | -0.5178 | 0.0475 | -0.4091 | 0.0172 |
| m$^1$I | -0.0913 | 0.0538 | -0.3149 | 0.0215 | -0.1344 | 0.0127 | -0.2375 | 0.0153 | -0.2616 | 0.0086 | -0.1457 | 0.0543 | -0.2332 | 0.0170 |

[a] Statistical significance was assessed using an unpaired two-sample $t$-test. Light red cells represent significance $p < 0.05$ and dark red cells represent significance $p < 0.01$.

[b] Standard deviation was calculated as transformed ($\log_2$) uncorrelated noncentral ratio distribution:

$$Log_2\sigma_z = \frac{1}{\ln(2)}\sqrt{\frac{\sigma_x^2}{\mu_x^2} + \frac{\sigma_y^2}{\mu_y^2}}$$

where $\sigma_z$ indicates the standard deviation of fold change, $\sigma_x$ and $\sigma_y$ indicate the standard deviation of measured peak heights of nucleoside modification levels in stressed and control conditions, respectively, $\mu_x$ and $\mu_y$ denotes the average peak heights of nucleoside modification levels in stressed and control conditions, respectively.

Table S3. Quantitative assessment of abundance-driven tRNA modification regulation by modification deviation ($MD_m$) scores[a,b].

| | 12 h | | | | 24 h | | | |
|---|---|---|---|---|---|---|---|---|
| Modification/condition | pH 7 | Heat (39 °C) | 0.5 mM MMS | 3 mM paraquat | pH 7 | Heat (39 °C) | 0.5 mM MMS | 3 mM paraquat |
| Am | 0.2117 | 0.3669 | 0.2564 | 0.2467 | 0.7254 | 0.4479 | 0.8231* | 0.5565 |
| Cm | 0.0545 | 0.0775 | 0.1989 | 0.3773 | 0.6110* | 0.5449* | 0.4852* | 0.4268* |
| Gm | 0.0208 | 0.1283 | 0.2158 | 0.1331 | 0.4360* | 0.4206 | 0.1919* | 0.1540 |
| I | 0.0167 | 0.1846 | 0.0639 | 0.1268 | 0.3197 | 0.3177 | 0.6258 | 0.5008 |
| $i^6A$ | 0.1053 | 0.0688 | 0.4474 | 0.1000 | 0.2198 | 0.2975* | 0.0579 | 0.2679* |
| $m^1A$ | 0.0566 | 0.0900 | 0.0951 | 0.0800 | 0.3395* | 0.3071 | 0.3593 | 0.2767 |
| $m^1G$ | 0.1633 | 0.1870 | 0.0518 | 0.0173 | 0.2007 | 0.2604 | 0.5419 | 0.2831 |
| $m^1I$ | 0.0807 | 0.0574 | 0.3271* | 0.1313 | 0.1302 | 0.1156 | 0.6677 | 0.4896 |
| $m^{2,2}G$ | 0.0013 | 0.0214 | 0.1067 | 0.2337 | 0.2750 | 0.3082 | 0.4388* | 0.2676 |
| $m^2G$ | 0.1280 | 0.1831 | 0.2044 | 0.2292 | 0.3346 | 0.3107 | 0.3974 | 0.3124 |
| $m^3C$ | 0.5089 | 0.3479 | 0.8233 | 0.4423 | 0.3112 | 0.0616 | 0.1187 | 0.0518 |
| $m^5C$ | 0.1094 | 0.0036 | 0.1731 | 0.1838 | 0.3270 | 0.3876 | 0.3619* | 0.3349 |
| $m^5U$ | 0.0567 | 0.2322 | 0.0432 | 0.0212 | 0.2509 | 0.365 | 0.2939 | 0.2495 |
| $m^7G$ | 0.0163 | 0.0347 | 0.1011 | 0.1317 | 0.3034 | 0.4705 | 0.3930 | 0.3533 |
| $mcm^5s^2U$ | 0.6029 | - | 0.0318* | - | 0.6080* | - | 0.9110* | - |
| $mcm^5U$ | 0.2242 | 0.8200* | 0.3981 | 0.8001* | 0.0261 | 1.0002* | 0.9670 | 0.9994* |
| $t^6A$ | 0.4583 | 0.1073 | 0.0840 | 0.9434* | 0.0672 | 0.0755 | 0.5556 | 0.2372 |

[a] Yellow highlight – $MD_m$ > 0.2; red highlight – $MD_m$ > 0.8

[b] Asterisks (*) mark values corresponding to modifications that are statistically significant (Fig. 3B).

Table S4. DNA oligonucleotide probes used for northern blotting.

| **Oligonucleotides** | Sequence (5'->3') | Source |
|---|---|---|
| SCer_tRNA-Glu(UUC) probe | CGATACGGGGAGTCGAACCCCGGTCTCCACG | This paper |
| SCer_tRNA-Gln(UUG) probe | GTCCTACCCGGATTCGAACCGGGGTTGTCCGG | This paper |
| SCer_tRNA-Lys(UUU) probe | GCCGAACGCTCTACCAACTGAGCTAACAAGG | This paper |
